# Primate V2 Receptive Fields Derived from Anatomically Identified Large-Scale V1 Inputs

**DOI:** 10.1101/2024.03.22.586002

**Authors:** Mahlega S Hassanpour, Sam Merlin, Frederick Federer, Qasim Zaidi, Alessandra Angelucci

## Abstract

In the primate visual system, visual object recognition involves a series of cortical areas arranged hierarchically along the ventral visual pathway. As information flows through this hierarchy, neurons become progressively tuned to more complex image features. The circuit mechanisms and computations underlying the increasing complexity of these receptive fields (RFs) remain unidentified. To understand how this complexity emerges in the secondary visual area (V2), we investigated the functional organization of inputs from the primary visual cortex (V1) to V2 by combining retrograde anatomical tracing of these inputs with functional imaging of feature maps in macaque monkey V1 and V2. We found that V1 neurons sending inputs to single V2 orientation columns have a broad range of preferred orientations, but are strongly biased towards the orientation represented at the injected V2 site. For each V2 site, we then constructed a feedforward model based on the linear combination of its anatomically-identified large-scale V1 inputs, and studied the response proprieties of the generated V2 RFs. We found that V2 RFs derived from the linear feedforward model were either elongated versions of V1 filters or had spatially complex structures. These modeled RFs predicted V2 neuron responses to oriented grating stimuli with high accuracy. Remarkably, this simple model also explained the greater selectivity to naturalistic textures of V2 cells compared to their V1 input cells. Our results demonstrate that simple linear combinations of feedforward inputs can account for the orientation selectivity and texture sensitivity of V2 RFs.

## INTRODUCTION

In the primate visual cortex, object recognition occurs via a series of transformations through hierarchically organized areas in the ventral visual pathway which originates in the primary visual cortex (V1) and terminates in the inferotemporal (IT) cortex, via intermediate areas V2 and V4^1^. As information flows through this pathway, neuronal receptive fields (RFs) become progressively larger and tuned to more complex image features. At the first cortical stage of this pathway, cells are tuned to simple image features such as the orientation of line segments^2, 3^, but as information is processed and passed along to higher stages, cells become selectively tuned to specific complex objects such as faces or hands^4–6^. The circuits, mechanisms, and computations that lead to the increased complexity of RFs along the cortical hierarchy have not been adequately characterized, even at the earliest transformation from V1 to V2. Computational models have attempted to understand how the more complex RF structure at one processing stage is derived from the preceding processing stage, but they have been based on measurements of isolated RFs instead of empirically identified anatomical and functional connections between cortical areas.

V2 is the largest of the primate extrastriate visual areas, it receives the vast majority of its cortico-cortical feedforward (FF) inputs from V1^7–9^, and its responses are abolished when V1 is silenced^10^. Like cells in V1, V2 neurons are selective for stimulus orientation and spatial frequency, but they can also be selective for more complex contours, such as elongated edges, angles and curves^11, 12^. In addition, V2 neurons demonstrate sensitivity to surface properties and selectively respond to naturalistic textures^13, 14^, texture and object borders, and stereoscopic depth cues ^15, 16^. V2 neurons have larger RFs compared to V1 neurons at comparable eccentricity, and exhibit greater contrast sensitivity^17, 18^. FF models of the visual system posit that the more complex RF properties of V2 neurons, such as their response to elongated or more complex contours, arise from pooling of inputs from V1 neurons with RF positions spread across visual space. Despite significant advances in our understanding of the anatomy and physiology of these areas, and despite hierarchical FF models of the visual system being central to many theories of visual object recognition^19–21^, how cells in V2 integrate inputs from V1 and how this integration accounts for the more complex properties of V2 RFs has not yet been demonstrated experimentally.

Our understanding of how V2 neurons encode information during natural vision is primarily based on theoretical studies, which have trained models to replicate some of the known response properties of V2 neurons^22–25^, or have generated data-driven models that rely on statistical analyses of responses of V2 neurons to natural stimuli^13, 26^. A major limitation of these theoretical studies is that their assumptions have not been tested physiologically and anatomically. For example, some of these models assume that V2 neurons use a sparse coding strategy and integrate inputs from a fixed number of V1 neurons, while other models are based on probabilistic representations of connectivity patterns between areas. Data-driven models, on the other hand, fit the data to mathematical models that can be difficult to interpret biologically. More importantly, none of these models have been constrained by realistic anatomical data and functional connectivity between cortical areas.

As the processing of visual contours and textures is believed to rely heavily on the computation of local orientations and their spatial relationships, in this study we focused on investigating the orientation and spatial organization of V1 inputs to V2. First, we combined functional imaging of orientation and retinotopic maps in V1 and V2 with anatomical labeling of V1-to-V2 inputs by injections of retrograde tracers into single V2 orientation columns. We then developed a computational model constrained by the functional connectivity of V1-to-V2 inputs identified in the first part of the study to investigate combinatorial rules and emerging functional representations in V2. We demonstrate that a simple FF model based on the linear combination of anatomically-identified V1 inputs to singe V2 orientation columns is capable of accurately reproducing the orientation tuning properties of V2 RFs. Consistent with previous investigations of V2 RFs, the V2 RFs derived by this model could be roughly classified as elongated V1-like filters, and filters with relatively complex spatial structures. Moreover, when applied to naturalistic texture images, the derived V2 filters showed higher sensitivity to the statistical dependencies in natural textures compared to V1 input cells, consistent with published experimental results^13, 14^. Our results demonstrate that simple FF mechanisms can account for the orientation selectivity and texture sensitivity of V2 RFs.

## RESULTS

### Feedforward inputs are orientation-biased and form complex spatial patterns

To understand the orientation and retinotopic organization of V1 inputs to V2, we obtained functional maps of orientation preference and retinotopy in these two areas, and visualized the V2 stripe compartments, by performing *in vivo* intrinsic signal optical imaging of V1 and V2 in macaque monkeys (**Fig. 1A-I**). We used these functional maps as a guide to target injections of retrograde neuroanatomical tracers (CTB-alexas tagged with different fluorophores, n= 10 injections in 4 animals) to V2 single orientation domains within the thick and pale cytochrome oxidase (CO) stripes, which, unlike thin stripes, contain well defined orientation maps (**Extended Data** Fig. 1D-E)^27, 28^. Following a post-injection survival period, animals were perfused with fixative, and the brain sectioned (see Methods). Fluorescent label in tissue sections was imaged on a confocal microscope (**Fig. 1F-G)** and microscopy images were aligned to the *in vivo* functional maps using the surface vasculature (**Extended Data** Fig. 1). For each labeled cell in V1 and pixel at the V2 injected sites we extracted a preferred orientation (PO) based on their location on the orientation maps (**Fig. 1E,K).** The relative positions of labeled V1 cells in visual space were determined based on their location on the retinotopic maps, using a cortex-to-visual field mapping procedure (**Fig. 1 H-K**). First, an area that encompassed the entire labeled field was outlined (yellow contour in **Fig. 1 H-J**), and its size was expressed in degrees of visual angle based on the number of retinotopic stripes it encompassed (for details see Methods). This area was then subdivided into a finely-tuned, uniformly distributed grid through an Elliptic Grid Generation approach^29^ (detailed in the Methods and Supplementary Methods). Grids initially contained 40000-360000 nodes, but were then re-sampled to give 0.02° resolution (**Fig. 1J**). Finally, each V1 cell was assigned to a retinotopic position in visual space based on its closest proximity to a given grid point (**Fig. 1K**). The map shown in **Fig. 1K** shows for each labeled V1 cell its retinotopic location and PO determined as described above.

**Figure 1.**
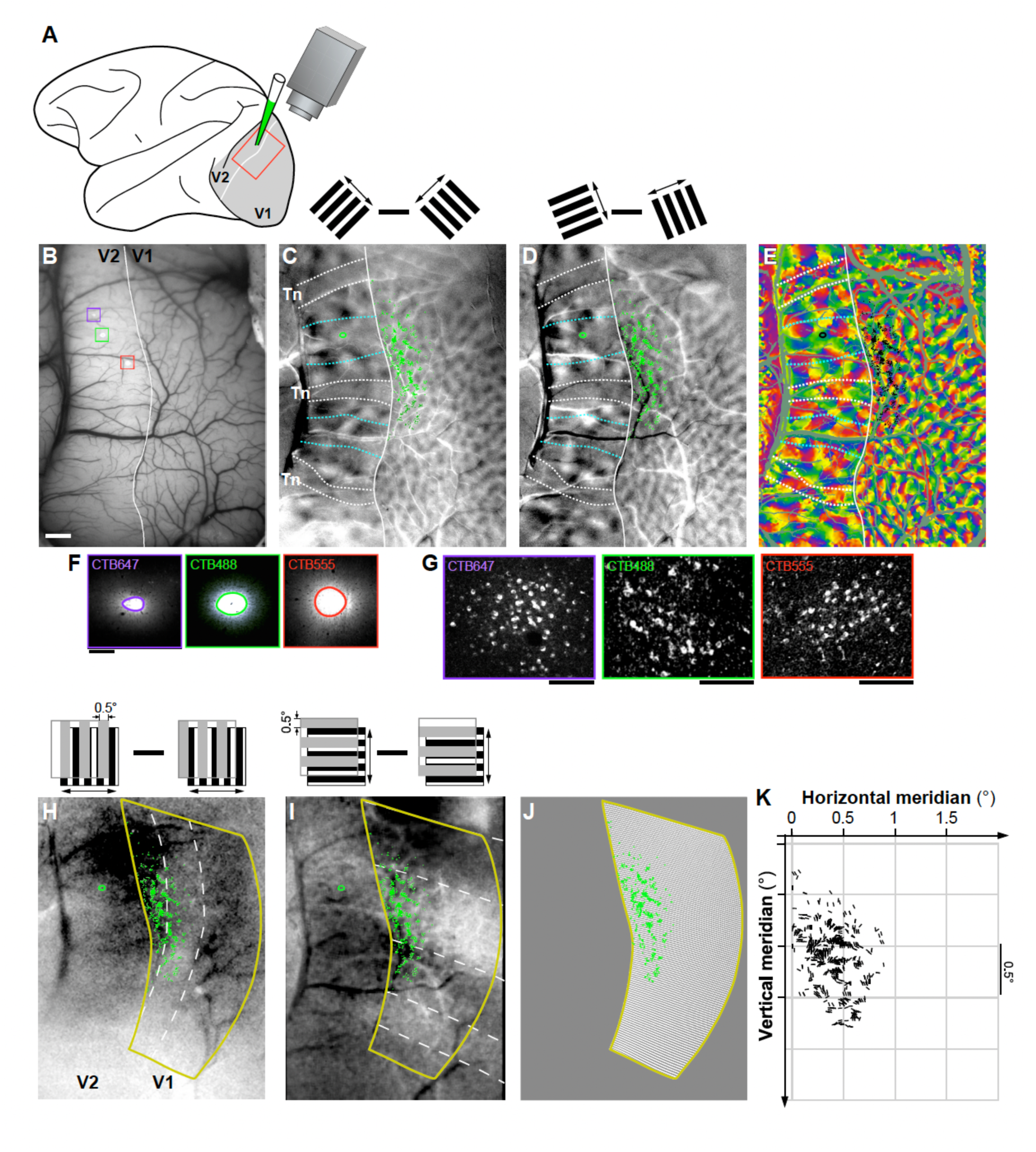
Experimental approach used to map in visual space the orientation and retinotopic organization of V1 inputs to single V2 orientation columns: example case MK373. **(A)** Schematics of the experimental approach. Intrinsic signal optical imaging (OI) of a region encompassing V1 and V2 (*red box*) was performed *in vivo* to obtain functional maps of stimulus orientation and retinotopy, and injections of retrograde tracers were made into a single orientation column in V2. (B) *In vivo* image of the surface vasculature, obtained under green light illumination, used as reference to position pipettes for tracer injections to specific functional domains, as well as to register functional maps to confocal images of histological sections (see **Extended Data** Fig. 1). In this representative case, three different tracer injections were made; confocal images of these injection sites are shown superimposed to the surface vasculature inside the *colored boxes*, and at higher magnification in panel (F): CTB647 (*purple*), CTB488 (*green*), and CTB555 (*red*). The *white contour* here and in (C-E) delineates the V1/V2 border based on the functional maps. Scale bar: 1 mm and valid for B-E and H-J. **(C, D)** Difference orientation maps obtained by subtracting responses to two orthogonally-oriented gratings (45°–135° and 22.5° – 112.5 °, respectively), as shown in the *insets* above the panels. The orientation maps in V2 show larger orientation domains than in V1, and a stripy organization, with regions having strong orientation responses corresponding to the thick (delineated by the *cyan dotted* contours) and pale stripes, and regions with weak or no orientation responses corresponding to the thin stripes (*Tn*; delineated by the *white dotted contours*). The outlines of the stripes shown on these maps are based on both the orientation and CO maps (see also **Extended Data** Fig. 1). Superimposed to the maps in (C-E) and (H-I) are manual plots of the locations of the V1 cells (*green dots* in C-D and H-I, *black dots* in E) labeled by the CTB488 injection, and the contour of the CTB488 injection site *(green* oval in V2). V1 label from the other tracer injections is shown **in Figure 2 and Extended data Fig. 2**. **(E)** V1 cells (*black dots*) labeled by the CTB488 injection (*black oval*) are shown superimposed to the composite orientation map. Other conventions are as in (C-D). **(F)** Confocal images of injection sites taken under 647 nm (left), 488 nm (middle), and 555 nm (right) light illumination. Injection sites in V2 were outlined manually as indicated by the *colored ovals*. Scale bar: 200µm. **(G)** Confocal images of V1 cells labeled by each respective tracer injection site taken under illumination with different light wavelengths. Scale bars: 100 µm. **(H,I)** Retinotopic maps generated by subtracting responses to 90° (H) or 0° (I) oriented gratings occupying complementary and adjacent strips (0.5° in width) of visual space (as shown in the *insets* above). This visual stimulation paradigm generates response stripes (manually delineated by *white dashed contours*) corresponding to the stimulated visual locations between the masks. The area encircled by the *yellow contour* is estimated to correspond to ∼2.5° along an axis parallel to the vertical meridian (corresponding to the location of the V1/V2 border) and 1.5° along the orthogonal axis. **(J)** Visual cortex-to-visual space mapping grid generated within the retinotopic mask (area delimited by the *yellow contour)*. This grid contains 125 points along the vertical meridian and 75 points along the orthogonal axis, which provides a resolution of 0.02° in both directions. CTB488-labeled V1 cells (*green dots)* are superimposed on the grid. For the purpose of the final analysis, the V1 cells that lay on vessels were discarded and are not shown here. **(K)** Visual field map of the retinotopic location and orientation preference of the CTB488-labeled V1 cells. The location of the vertical meridian is at 0° on the Horizontal Meridian axis and corresponds to the location of the V1/V2 border on the brain. As there is no landmark for the horizontal meridian representation on the brain, its location relative to the labeled cells could not be determined accurately, and as such it is not indicated on the vertical meridian axis. The labeled field is located at parafoveal eccentricities (3-7°).

Figure 2 presents results from 4 representative V2 injection cases; the remainder of cases (n=6) are shown in **Extended Data** Figure 2. Tracer injection sites in V2 (ranging in diameter between 200 µm and 580 µm) involved mostly layers 2-4, and were mostly confined to one or two V2 orientation columns. Depending on their size, a single injection site labeled between 162 and 7402 V1 neurons in layers (L) 2/3. Retrograde label was also found in L4A-B. However, our analysis focused solely on V1 inputs from L2/3, as these neurons are the ones contributing to the functional responses recorded with optical imaging from the cortical surface.

**Figure 2.**
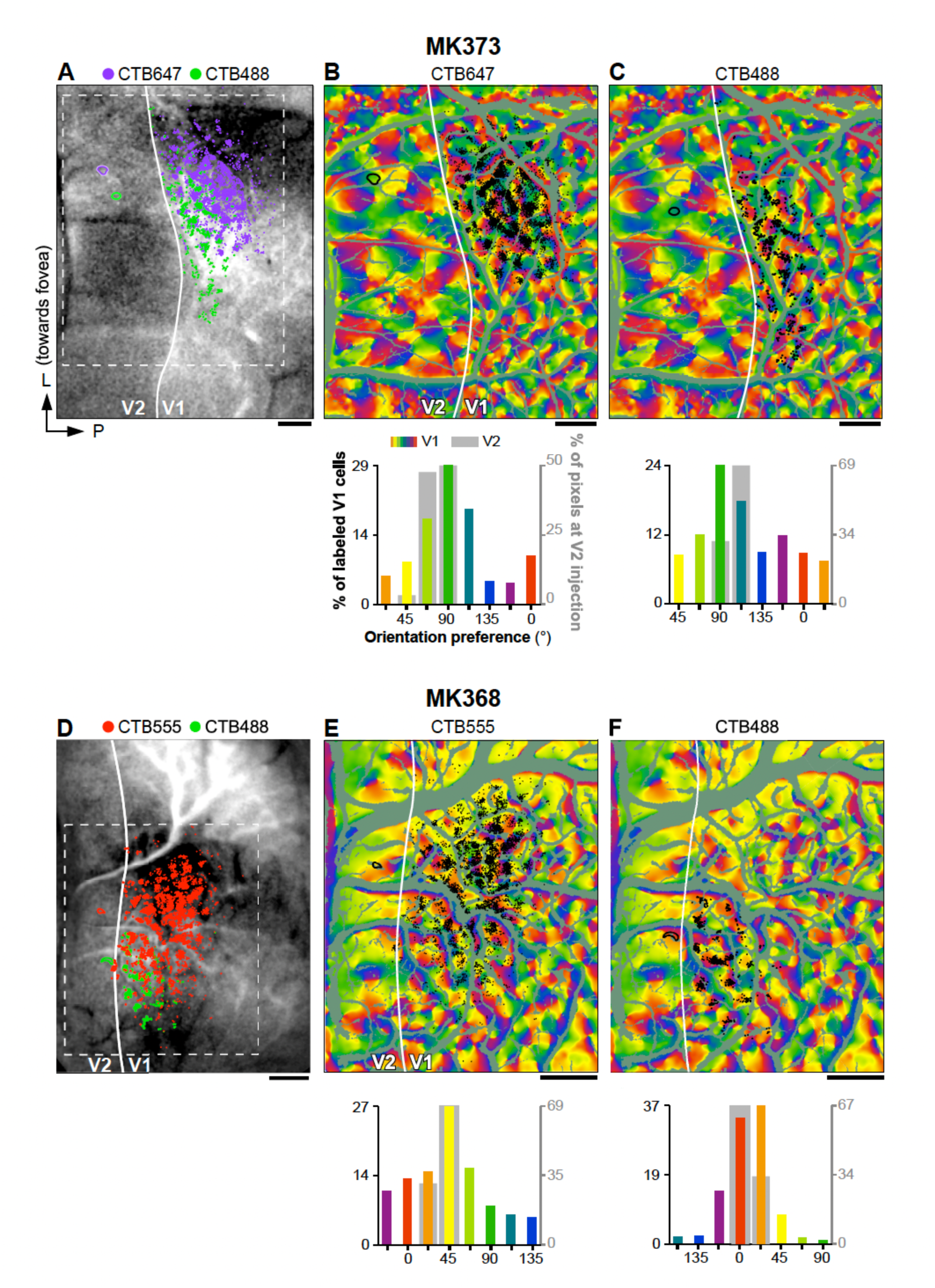
Orientation organization of V1 inputs to V2 orientation columns in 4 representative cases. **(A)** CTB647 (*purple oval*) and CTB488 (*green oval*) injection sites in V2, and resulting labeled cells in V1 (*purple and green dots*, respectively) are shown superimposed onto the imaged 0° retinotopic map for case MK373. The orientation map for the region inside the *dashed white box* is shown in panels (B-C). Other conventions are as in **Fig. 1. (B)** TOP: The locations of the CTB647 injection site in V2 (*black oval*) and resulting labeled cells (*black dots*) in V1 are shown superimposed to the composite orientation map for the same case shown in (A). BOTTOM: Distribution of POs at the V2 CTB647 injection site (*gray bars*; right Y axis) and under the CTB647 labeled cells in V1 (*colored bars*: left Y axis). **(C)** Same as (B), but for injection case MK373-CTB488. **(D-F)** Same as (A-C), but for two additional injection cases: MK368-CTB555 (D-E), and MK368-CTB488 (D,F). Scale bars: 1mm. The reminder of cases used in this study are shown in **Extended Data. Fig. 2**.

V1 L2/3 neurons labeled by a single injection had POs that were distributed broadly, but were strongly biased towards the POs represented at the injected V2 site (Fig. 2B**,C****,E,F,** and **Extended Data** Fig. 2A,B**,D-G**). Specifically, on average across the population of injection sites 47.7%±13.2% (SD) of labeled V1 cells had a PO within ±22.5° of that at the V2 injection site. However, there was variability among the different injection cases, with some injections showing narrower and other broader distributions of POs in the labeled V1 cells (range: 35.4%, for case MK365-CTB555, to 72.7% for case MK368-CTB488). There was a tendency for pale CO stripes to show narrower distributions than thick stripes, but differences among CO stripe types were not statistically significant (Pale-lateral: 53.6%±16.6, Pale-medial: 47.9%±13.8, thick 38.7%±4.7%). When mapped in visual field coordinates, the entire field of V1 input cells labeled by each V2 injection encompassed approximately 1-2° of visual space, i.e. slightly larger than the average RF of a V2 cell in parafoveal V2^30^ (Fig. 3 and **Extended Data** Fig. 3).

**Figure 3.**
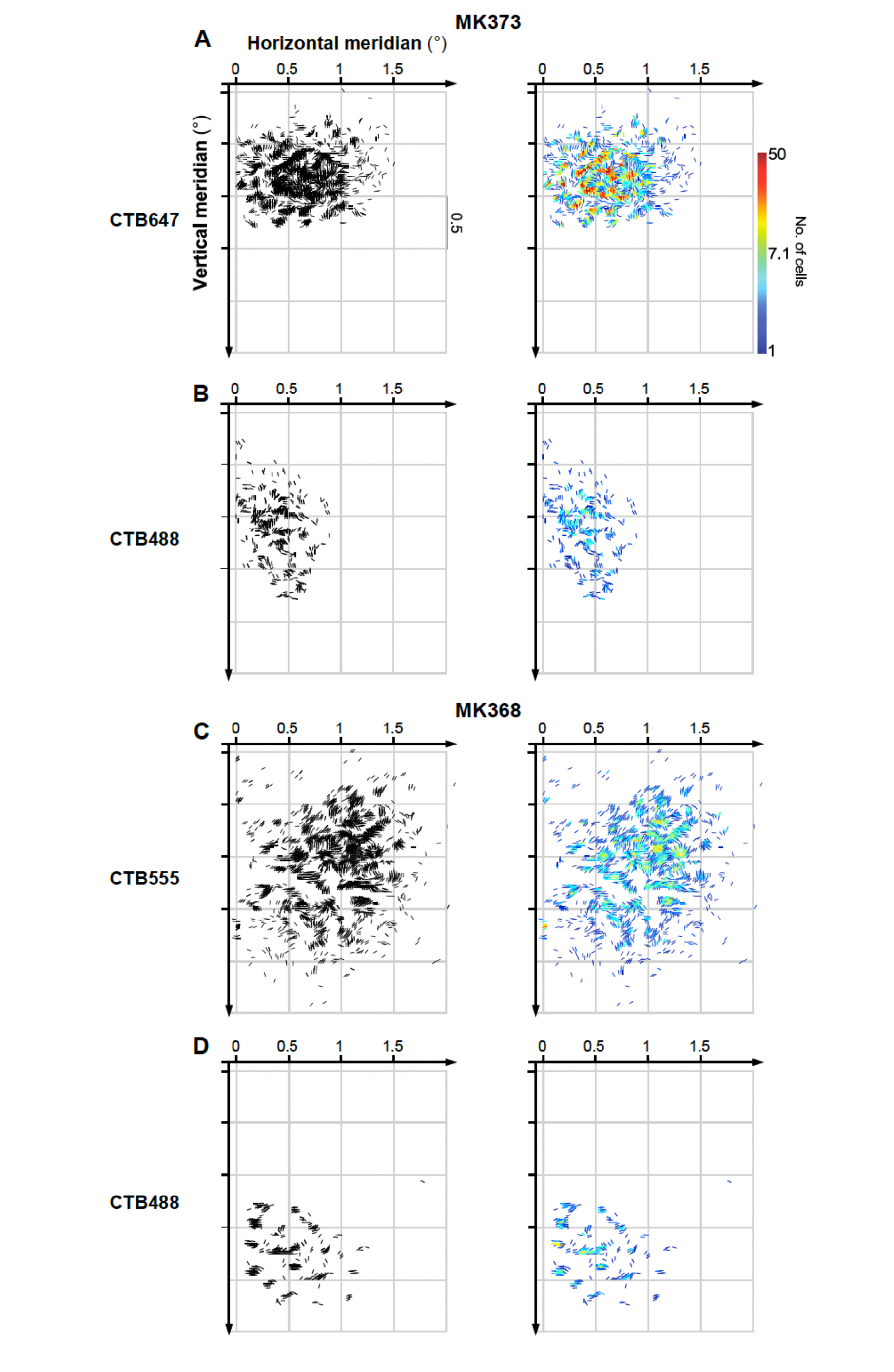
Visuotopic maps of V1 inputs to V2 orientation columns in 4 representative cases. **(A)** Visual field maps of POs and retinotopic layout of labeled V1 cells for case MK373-CTB647 (same case as in **Fig. 2A-B**). LEFT: Each labeled V1 cell is represented as a black oriented line segment centered on each cell’s RF location in visual space with the line orientation corresponding to the cell’s PO. RIGHT: the same map is shown as a color-coded version, in which the *colors of lines* indicate the number of labeled cells (in logarithmic scale) mapped at each cell’s RF location in visual space. **(B-D**) Same as in (A) but for different injection cases as indicated (same cases as in **Fig. 2**). Whereas relative cell density is better revealed by the color maps, the overall pattern of the V1 inputs is better captured by the black maps. The visuotopic maps for the reminder of cases are shown in **Extended Data** Fig. 3.

The data in Fig. 2 and **Extended Data** Fig. 2 demonstrate that each V2 injection labeled input neurons in a discrete region of V1 (for the example case MK373-CTB647, this corresponds to the area of the orientation map inside the white contour in **Extended Data** Fig. 4A). Moreover, within this labeled V1 region, labeled neurons were concentrated in patches, rather than being uniformly spread over the region. This raised the question of whether the labeled patches are sampling V1 orientations selectively, with a bias towards the PO of the V2 injection site, or non-selectively. To address this question, we performed three kinds of analyses. First, we asked whether the orientation bias observed in the distribution of POs of the V1 inputs simply reflects an overrepresentation of a subset of orientations in the V1 orientation map. To answer this question, for each injection case we compared the distribution of POs of the labeled V1 cells with the distribution of POs of all the pixels within the V1 labeled cell field (e.g. the area of the orientation map inside the white contour in **Extended Data** Fig. 4A), as well as within the entire imaged V1 field of view, excluding blood vessels (i.e. the entire map V1 region shown Fig. 1E). In contrast to the V1 input cells labeled by the V2 injections (Fig. 2B for the example case MK373-CTB647), the POs of all pixels within the labeled V1 field (**Extended Data** Fig. 4B **Top panel**) or within the entire V1 imaged field of view (**Extended Data** Fig. 4F **Top panel**) showed no bias towards representing the PO at the V2 injection site. The complete set of control data showed a slight bias towards multiple orientations, but the biased orientations differed in different cases. For example, in case MK373-CTB647, the orientation map within the labeled field and within the entire imaged field of view showed a slight overrepresentation of the cardinal axes (0, 90°; top panels in **Extended Data** Fig. 4B and F, respectively), but in other cases the orientation maps were slightly biased towards non-cardinal orientations. Importantly, the biased orientations in the control data did not match the bias in the real data (the V1 labeled cells), and a chi-square goodness of fit test, showed that the distributions of POs in the control and real data were significantly different from each other for all cases (p <<0.05, degree of freedom = 7, i.e. number of bins minus 1). These results suggest that V1-to-V2 connections arise from selective regions in the V1 map and that the bias in the distribution of POs in these inputs does not reflect an orientation bias intrinsic to the V1 orientation map.

We performed two additional statistical tests to determine whether the observed orientation bias in the distribution of POs of the labeled V1 cells could result from a random spatial pattern of V1-to-V2 connections within the V1 projection field, or from the observed spatial pattern of connections placed randomly within V1. To this goal, we simulated control data by two different random placement strategies. In the first analysis, the observed distribution of POs of the labeled V1 cells was compared to the distribution of POs under an equivalent number of pixels randomly selected within the labeled cell field 1,000 times (**Extended Data** Fig. 4A, B **bottom panel, C**). As a second test, simulated data were generated by shifting the real pattern of labeled cells (i.e. preserving the spatial pattern) to >1,000 different locations within the imaged field of view (**Extended Data** Fig. 4E, H), and then calculating the resulting distribution of POs (**Extended Data** Fig. 4F **bottom panel, and G**). For both analyses, in contrast to the observed V1 distribution, the simulated V1 distributions showed no bias towards representing the PO at the V2 injection site (bottom panels in **Extended Data** Fig. 4B, F). Given the cyclical nature of the orientation data, we applied circular statistics as a summary metrics for statistical comparison of observed and simulated distributions; specifically, we estimated the mean resultant length (MRL) and the circular standard deviation (CSD), as described by Fisher (1993)^31^. The circular statistics calculated from the observed distributions consistently fell at the extremes of the distributions generated by the simulations (**Extended Data** Fig. 4D, I, and **Extended Data Tables 1-2**). Further statistical analysis, utilizing the Kolmogorov–Smirnov test with a Bonferroni-corrected family-wise p-value of 0.05, rejected the hypothesis that the observed distributions were reproducible by random sampling from the imaged V1 orientation maps.

**Table 1.** Statistical Analysis of difference in texture sensitivity between V1 and V2 cells in the model.

| **Case** | **CO stripe** | **Mean MI – V1** | **Mean MI – V2** | **t-stats** | **SD** | **p-value** |
| --- | --- | --- | --- | --- | --- | --- |
| **MK373 – CTB 555** | Pale Medial | 0.164 | 0.180 | 2.33 | 0.058 | 0.021* |
| **MK373 – CTB 488** | Thick | 0.164 | 0.220 | 5.09 | 0.071 | 0.000* |
| **MK373 – CTB 647** | Pale Lateral | 0.164 | 0.190 | 2.34 | 0.065 | 0.020* |
| **MK368 – CTB 488** | Pale Lateral | 0.161 | 0.150 | -2.05 | 0.052 | 0.042* |
| **MK368 – CTB 555** | Pale Medial | 0.164 | 0.170 | 0.76 | 0.055 | 0.448 |
| **MK365– CTB 488** | Pale Medial | 0.160 | 0.170 | 0.90 | 0.059 | 0.370 |
| **MK365 – CTB 555** | Thick | 0.163 | 0.300 | 8.31 | 0.116 | 0.000* |
| **MK365 – CTB 647** | Pale Lateral | 0.163 | 0.250 | 6.89 | 0.089 | 0.000* |

SD: estimate of population standard deviation

These results suggest that V1-to-V2 connections sample V1 orientations selectively, and preferentially link neurons in these two areas having similar POs. Like-to-like connectivity could represent the anatomical substrate for the processing of elongated oriented contours and orientation-selectivity in V2 cells, but it would not lead to the more complex RF structures also described for V2 cells. In fact, the observed like-to-like connectivity is not absolute, as many V1 inputs contacted V2 regions having different (>±30°) and even orthogonal POs. Computer simulations further indicated that this “imperfect’ like-to-like connectivity was not due to the V2 injection site not being confined to single V2 orientation columns. Specifically, we simulated V1 PO distributions under conditions of perfect like-to-like connectivity, and then compared these to the real distributions. These simulations took into account that orientation responses had been measured with gratings separated by 22.5°orientation, that the computation of orientation preferences for V1 neurons included Gaussian smoothing, and that the V2 injection site was not perfectly confined to a single orientation column. We tested whether the observed variation in input POs was broader than simulated by these factors (see Methods for details), and found that the observed distributions in PO were significantly broader (p << 0.05) than the simulated distributions (**Extended Data** Fig. 5), underscoring the complexity of the real neural connections.

To gain greater insights onto the spatial distribution of POs of the labeled V1 input cells, for each case we plotted an oriented line element at each cell’s estimated location in visual space (as described in Fig. 1K). The orientation of these lines matched the PO for each corresponding cell (Fig. 3 and **Extended Data** Fig. 3**). In** Fig. 3 and **Extended Data** Fig. 3, for each injection case we show black and color-coded versions of the same visuotopic map. In both the black and color map, each cell is represented as an oriented line, and in the color map, the color scale indicates the number of cells at each retinotopic location. The resulting maps revealed complex patterns of orientation flows, such as collinear and parallel edge elements, angular and curvature configurations, and textural patterns. This complexity suggests that the integration of these local line elements by V2 cells could shape the more complex RF properties of V2 cells such as their selectivity for angles^12^, and textures^32^. This led to the important questions of how V2 cells combine information from these oriented V1 RFs, and what RF properties emerge from this combination. In the next section, we tested whether a simple feedforward model can provide adequate answers.

### The orientation tuning of V2 columns is predicted by a linear combination of their V1 inputs

To understand how V2 cells integrate their V1 inputs, and what RF characteristics emerge from this integration, we developed a simplified feedforward model. This model incorporated some simplifications that were dictated by the limitations of our data. As our imaging experiments did not characterize the phase sensitivity of the imaged V1 cells, we explored two model variants with V1 cells modeled as either complex or simple. First, V1 cells were modeled as phase-invariant (complex RFs), comprising the three layers illustrated in Fig. 4A. The first layer consisted of all the labeled V1 cells projecting to the V2 injected site. Each cell was represented by four Gabor filters with spatial phases offset by 90 degree (two with an even-symmetric spatial structure and two with an odd-symmetric spatial structure)^33^. The parameters of the Gabor functions, including orientation and aspect ratio of the Gaussian envelope, were estimated from our recorded functional imaging data by fitting tuning curves of a bank of Gabor filters to the tuning curves measured from the imaging maps (see Methods for details). The spatial frequency of the sinusoidal carrier was set at 1 cycle/ degree to correspond to the spatial frequency of the gratings used to measure orientation selectivity in our imaging experiments. The responses of V1 complex cells to an input image (visual stimulus) were modeled in the second layer as the sum of responses of the four Gabor filters after half-wave positive rectification^34^. Finally, in the third layer, a V2 cell’s response was modeled as a weighted spatial sum of the responses of its V1 input cells, so the V2 RF reflected both the retinotopic locations and the strength of connection of the V1 cells. In the simple cell model, instead, V1 cells were modeled as phase-sensitive simple cells (single Gabor filters in the same phase for all cells), and then the weighted spatial sum of the V1 responses was taken in the same way as for the complex cell model.

**Figure 4.**
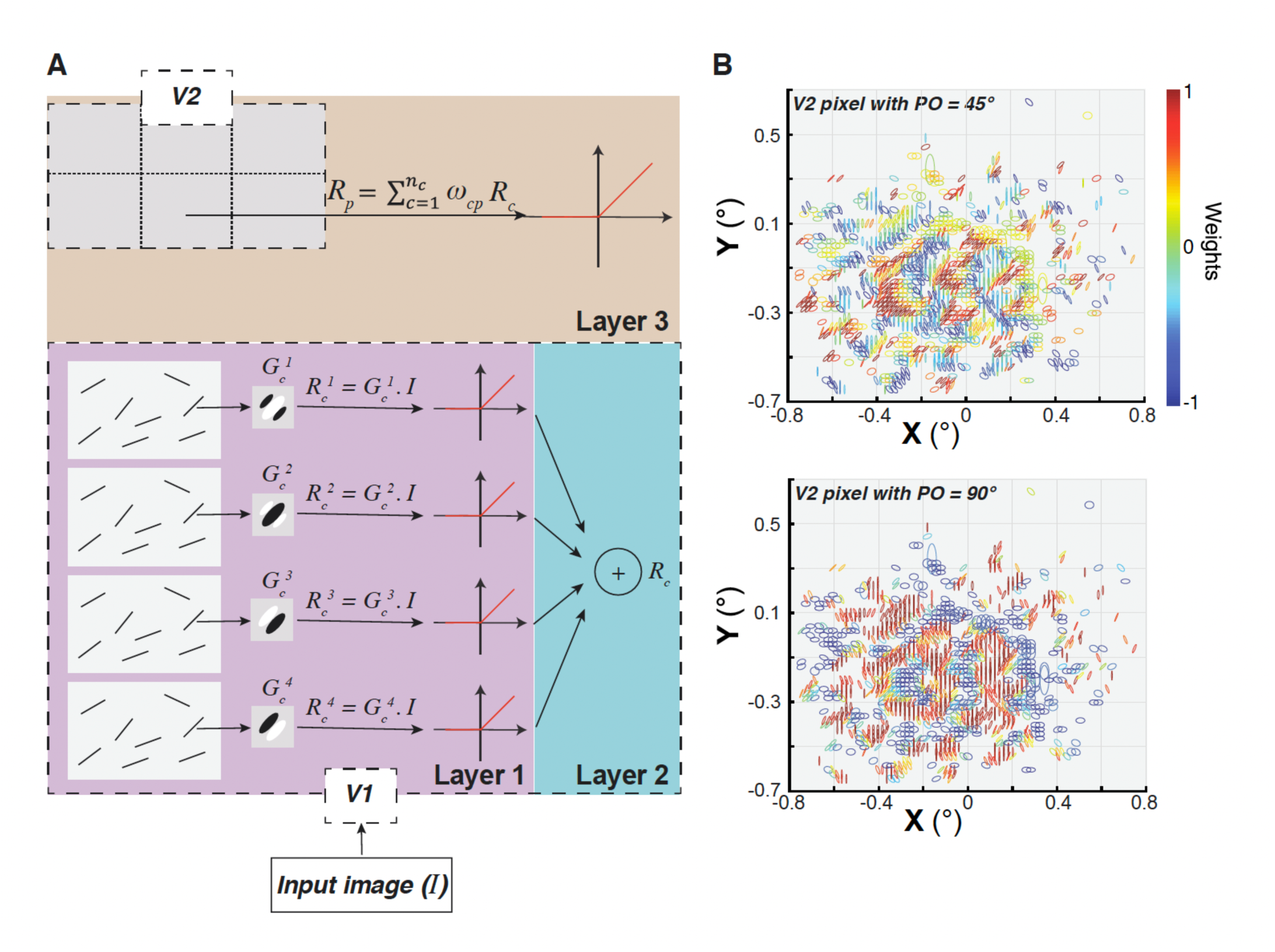
The linear complex-cell feedforward network model. **(A)** The key components of the three-layer complex-cell model are illustrated. **Layer 1** (*purple*): the receptive field (RF) of each V1 cell (*c*) is characterized by four two-dimensional Gabor functions (*G_c_^i^*,*i* = 1:4), each having a distinct spatial phase 0, 180, 90 and 270 degrees, respectively). The response of each model V1 cell’s RF to an input image, *I*, is quantified as the half-wave rectified dot product between the image and the corresponding Gabor function (*R_c_^i^* = *G_c_^i^*.*I*). **Layer 2** (*cyan*): a complex V1 cell’s response to the input image is modeled by summing responses of the four RFs modeled in layer 1. **Layer 3** (*khaki*): the response of a V2 pixel/cell to the input image, *R_p_*, is modeled as a weighted sum of all inputs from V1. *ω_cp_* represents the weight for a V1-V2 cell-pixel pair **(B)** Spatial distribution of various parameters employed in constructing the RFs of two distinct V2 cells/pixels in case MK373-CTB647. These cells/pixels exhibit POs of 45° (TOP) and 90° (BOTTOM). In these plots, each projecting V1 cell is depicted as an ellipse, aligning with the orientation and aspect ratio of its associated Gabor function. The *color*s used here denote the weights within the linear model.

V2 RFs calculated as the weighted sum of their V1 inputs would reflect the simplest combination rule. Since a regression would be ill-conditioned, because of rank deficiency and multicollinearity between similarly tuned V1 inputs, the weight for each V1 cell - V2 pixel pair was estimated as the dot product of their mean-subtracted and normalized tuning curves (see Methods for details). Figure 4B shows the parameters of the Gabor functions, including orientation, location and aspect ratio, along with the weights used for modeling two example V2 cells/pixels. The largest weights are, as expected, from V1 inputs with similar PO to the V2 cell, and the figure shows that such inputs are distributed in small clusters over the whole V2 RF, and are flanked by clusters of V1 inputs with orthogonal POs having negative weights. This organization is reminiscent of previous analyses of responses of V2 neurons to natural stimuli, showing that local excitatory edges have nearby suppressive edges with orthogonal orientations^26^; this organization increases the sparseness of responses to natural images and enhances the local representation of excitatory signals.

To picture the spatial configurations of V2 RFs resulting from the linear combination, the RFs of V1 cells were spatially summed using the weights from the linear model, separately for even-symmetric and odd-symmetric filters. The resulting RFs (Fig. 5) could be broadly categorized into two types: elongated filters resembling V1 RFs with distinct ON and OFF oriented regions, but usually more elongated than typical V1 cells (e.g. cell MK368-CTB488 in Fig. 5), and more complex RFs containing several non-oriented regions (e.g. cell MK373-CTB647 in Fig. 5) as well as multiple oriented regions (e.g. cells MK365-CTB488 odd, MK373-CTB555 and CTB488, MK368-CTB555 in Fig. 5), akin to those reported previously for experimentally measured RFs of macaque V2 cells^30^. In some cases, the RFs that are the sums of odd filters differ from the RFs that are the sum of even filters by just a luminance polarity reversal, but in other cases the difference between these RFs is more marked. Note that results are shown for only 8 injection cases because for two cases (MK335-CTB488 and CTB55) we lacked a complete set of orientation responses (in these two cases, 8 orientations were sampled in four separate trials as opposed to a single trial which would allow appropriate baseline correction for tuning curve analysis which is essential step in model estimation).

**Figure 5.**
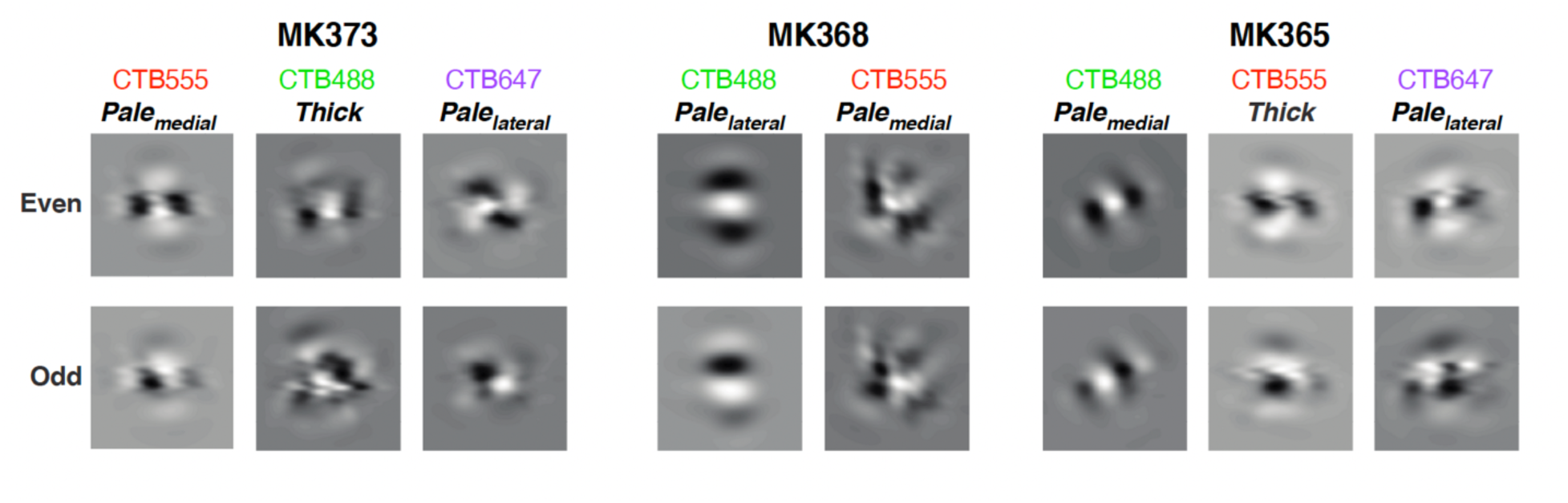
Spatial structure of modeled V2 receptive fields. Representative examples of modeled V2 RFs in 8 different injection cases, constructed by combining odd- or even-symmetric V1 Gabor filters. Both the tracer and CO V2 stripes injected are indicated for each case.

The combination weights were not collectively optimized to fit any V2 properties, and the dot products were largest for the V1 cells whose orientation tuning is closest to the orienation tuning of the V2 target, so we evaluated how well the calculated linear combination predicts the responses of V2 cells/pixels to oriented gratings, i.e. their orientation preference and tuning. To accomplish this, we conducted an eight-fold cross-validation analysis. In this analysis, the model was iteratively constructed eight times, each time utilizing the responses of V1 cells and V2 pixels to seven of the eight orientation stimuli. The experimental data corresponding to the omitted orientations served as the validation set. Figure 6A displays the predicted (using the complex cell model) versus observed responses of V2 cells/pixels to the excluded grating stimuli. The prediction error was quantified as a relative error (defined as the absolute difference between predicted and measured responses of V2 cells/pixels to grating stimuli, divided by the measured response range across all 8 orientations), averaged across eight iterations and then across all V2 cells/pixels for each case (Fig. 6B). The complex cell model exhibited robust performance across all eight injection cases (Fig. 6B) with a grand average relative cross-validation error of 0.16±0.04 (standard deviation across all 8 cases). For the simple cell model, the relative cross-validation error for the majority of injection cases was higher compared to the complex cell model (**Extended Data** Fig. 6A-D), with mean relative cross-validation errors of 0.33±0.15 (for the model based on odd-symmetric V1 filters) and 0.26±0.13 (for the model based on even-symmetric V1 filters). Overall, both linear feedforward models (simple and complex) demonstrated strong predictive capability for the orientation preference of V2 cells/pixels with a median absolute error (defined as absolute value of predicted PO minus measured PO) of about 5° for the complex cell model, and of about 5° and 6° for the odd and even simple cell model, respectively (Fig. 6C **and Extended Data** Fig. 6E-F). However, the complex cell model performed noticeably better than the simple cell model in predicting the orientation tuning width of the modeled V2 cells (Fig. 6D and **Extended Data** Fig. 6G-H) (median absolute error for HWHH = 5° versus 37°, respectively). In subsequent sections we will be focusing on the complex cell model, as the latter performed better overall than the simple cell model.

**Figure 6.**
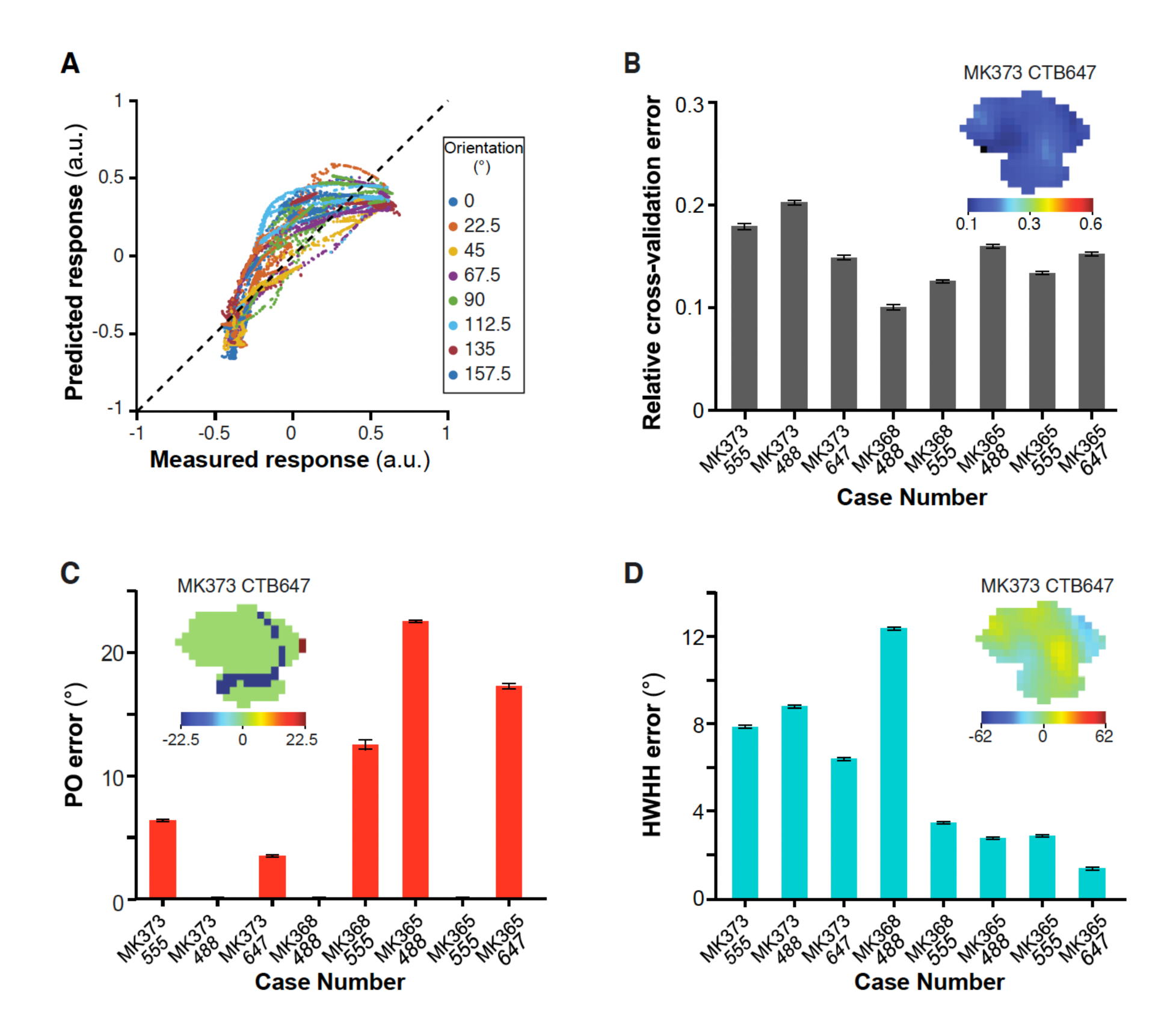
Performance of the complex-cell feedforward linear model. **(A)** Responses of V2 cells/pixels to excluded grating stimuli of 8 different orientations predicted by the complex cell model in the leave-one-out (LOO) procedure (y axis) versus responses measured experimentally (x axis). This plot shows data for all V2 cells/pixels in eight injection cases. The mean correlation coefficient between measured and predicted responses across the 8 LOO tests is 0.88±0.04. **(B)** Averaged relative cross-validation error for each injection case. Zero indicates a (perfect) prediction, while a value of 1 indicates that the difference between predicted and measured response equals the range (i.e. max minus min) of the cell’s measured response to the full set of 8 orientations; in practice the relative errors were never >0.6. The *inset* shows a color-coded relative error map at the V2 injection site for a representative case (MK373-CTB647). **(C)** Averaged absolute error (defined as the absolute value of predicted PO minus measured PO) in the model’s prediction of the preferred orientation (PO) of V2 cells/pixels. The *inset* is a color-coded signed error map calculated for each pixel at the V2 injection site for case MK373-CTB647. **(D)** Averaged absolute error in the model’s prediction of the width of the tuning curves (HWHH: half width at half height) of V2 cells/pixels. The *inset* is a color-coded signed error map calculated as predicted HWHH minus measured HWHH for each pixel at the V2 injection site for case MK373-CTB647 . Error bars: s.e.m.

### Texture sensitivity can emerge from linear feedforward connections

We asked what V2 RF response properties emerge from the simple linear combination of V1 inputs, compared to the responses of their V1 inputs. The elongation of some RFs and the complex organization of unoriented ON and OFF sub-regions of other V2 RFs shown in Figure 5 suggests that both may facilitate the representation of naturalistic visual textures in this cortical area^13^. To investigate this, we measured the responses of V1 and V2 model cells to a large set of synthesized naturalistic texture images, which include the higher-order statistical dependencies found in natural textures^35^, and to their spectrally-matched noise images, which lack these dependencies. For each of several original images of visual textures, 30 samples of naturalistic texture images and their noise counterparts were synthesized, forming what we refer to as a texture family (see Methods). We used 97 texture families, i.e. 97x30= 2,910 images. Images were cropped to be square shape, resized to 320 x 320 pixels, and masked using a circular mask of 3.2° in diameter (approximately twice the RF size of the model V2 cells), and were presented at the center of the V1 input cells aggregate RF. To maximize the response to the texture pattern of cells having different orientation preferences, each texture and noise image pair was presented at 8 different rotations. A texture modulation index (MI), defined as response to texture minus response to noise divided by the summed response (texture + noise)^13^, was calculated at each texture orientation and we analyzed results at the orientation that provided the most significant differential response between texture and noise images, i.e. the orientation that provided the largest mean over variance value). A higher MI corresponds to a model cell being more sensitive to the higher-order statistical dependencies that are shared by different samples of a naturalistic texture. We found that, compared to model V1 cells, the model V2 cells exhibited stronger responses to subsets of texture families relative to their noise counterparts, as indicated by higher MI values . Specifically, Figure 7A shows for each case the mean MI value of V2 vs V1 cells (averaged across cells) for each of 97 texture families. For 6 out of 8 cases, average mean V2 MI (across all 97 texture families) was higher than average mean V1 MI. The average mean V2 MIs for the entire population was 0.19±0.04 (S.D.), while it was 0.16±0.01 for V1 (yellow dot in Fig. 7A). Figure 7B shows for the entire population of V1 (top) and V2 (bottom) cells across all 8 cases the distribution of MIs averaged across all texture families for each cell. V2 cells showed a greater spread of MI values than V1 (range for V2 0.14-0.43, for V1 0.15-0.17), and a higher average mean MI value (V2: 0.19±0.1.; V1: 0.16±0.06). Naturalistic textures vary considerably in their texture structure, and the closer the RF structure is to the texture structure, the higher will be the MI for that texture-family. MI averaged over all texture families may, thus, underestimate the ability of V2 cells with complex RFs to differentiate between selected naturalistic textures and their matched noise. In keeping with this conjecture, the difference in mean MI values between V1 (0.33±0.03) and V2 (0.48±0.16) became even more pronounced when only the maximum MI value was taken for each cell (Figure 7C). The histograms in Fig. 7C show that the mean MI for V2 is larger than all the MI for V1, so all values larger than the mean are exclusively from V2 model cells. Statistical analysis (t-test, p< 0.05) demonstrated a significant difference in mean MI values between V1 and V2 across all texture families in 6 out of 8 injection cases (**Table 1**). Notably, this distinction was particularly pronounced for the thick (t-test, p < 10^-11^) and pale-lateral (t-test, p < 0.001) CO stripe types (**Extended Data** Fig. 7) compared to the pale medial-stripes.

**Figure 7.**
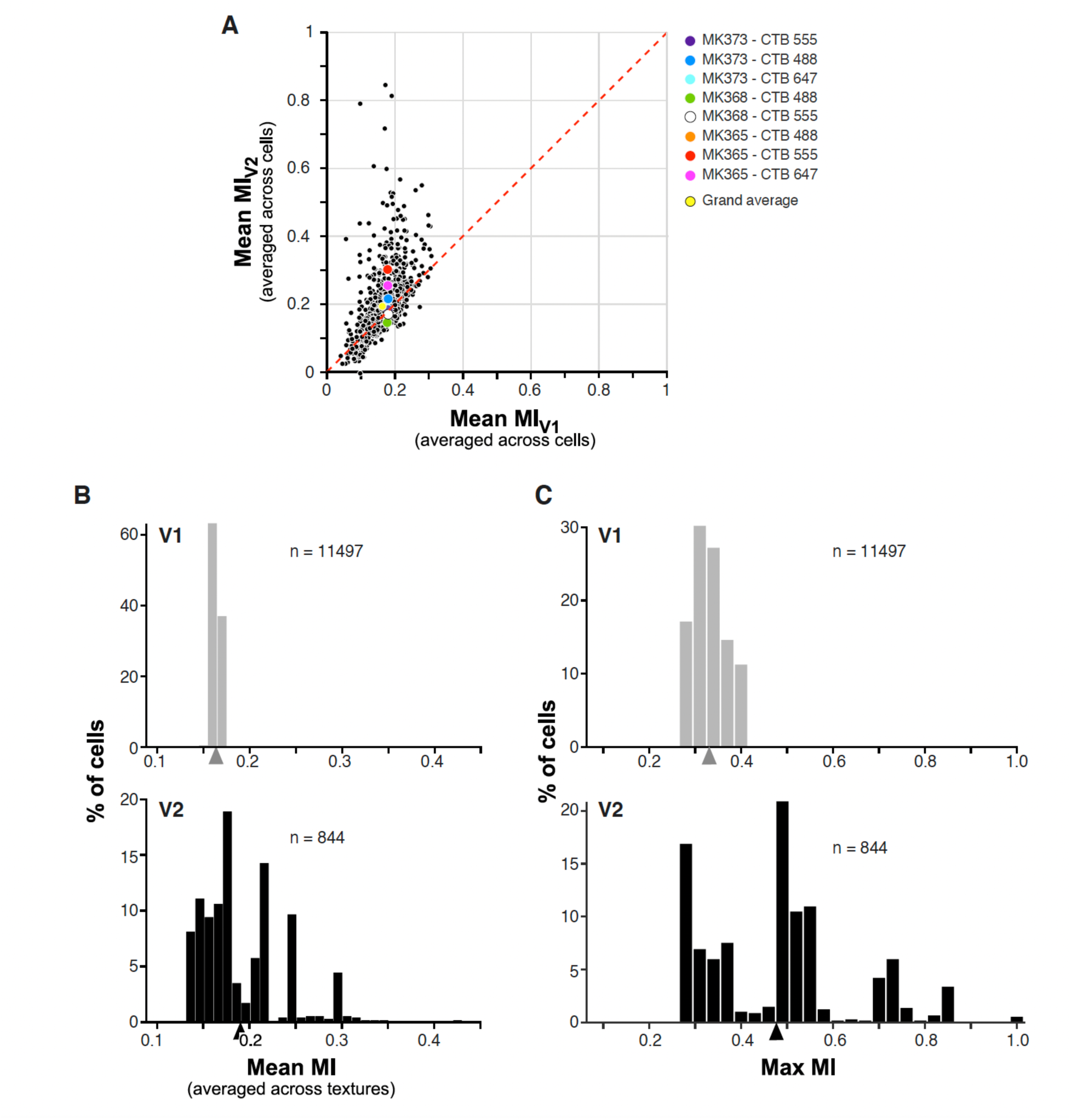
Responses of model cells to naturalistic textures. **(A)** Each *black dot* in the scatter plot represents the MI for a given texture family averaged across all modeled V2 and V1 cells for each injection case; there are a total of 97 dots per case, corresponding to the number of texture families used. *Colored dots* indicate mean MIs across all 97 texture families for each case. *Yellow dot* is the grand average across the entire population of cells. **(B)** Distribution of mean MIs for V1 (TOP) and V2 (BOTTOM) model cells. Here, for each V2 and V1 cell we calculated the average MI across all 97 texture families. *Arrowheads*: population mean. **(C)** Distribution of MIs for V1 (top) and V2 (Bottom) model cells. Here, for each cell we took the MI with the largest value.

In Figure 8A we show representative images taken from the 15 texture families that evoked the highest responses from model V2 cells ranked by V2 MI values. MI values in response to these textures were higher in V2 than in V1. The converse was true for the 15 texture families that evoked the lowest responses from V2 cells, most of which instead evoked higher MI values in V1 than in V2 (Figure 8B). For comparison, Figure 8C shows the 15 texture families that evoked the highest responses from V1 cells ranked by MI value. Notably, V1 and V2 cells preferred many of the same texture families, but MI values in response to these preferred textures were generally higher in V2 than in V1. Moreover, V2 responses to these textures were much more robust, i.e. less variable than V1 responses to the same textures, as indicated by the much larger error bars for V1 mean MIs compared to those for V2 MIs. The average standard errors for the MIs across all textures were 0.164, and 0.002 for V1 and V2, respectively, and the mean divided by the standard error was 112.2 and 1.1 for V1 and V2, respectively. These results along with the statistics in Fig. 7 demonstrate that even simple linear feedforward connections from identified V1 inputs to target V2 cells can generate the increased selectivity to naturalistic textures that has been claimed to be a signature feature of area V2^32^.

**Figure 8.**
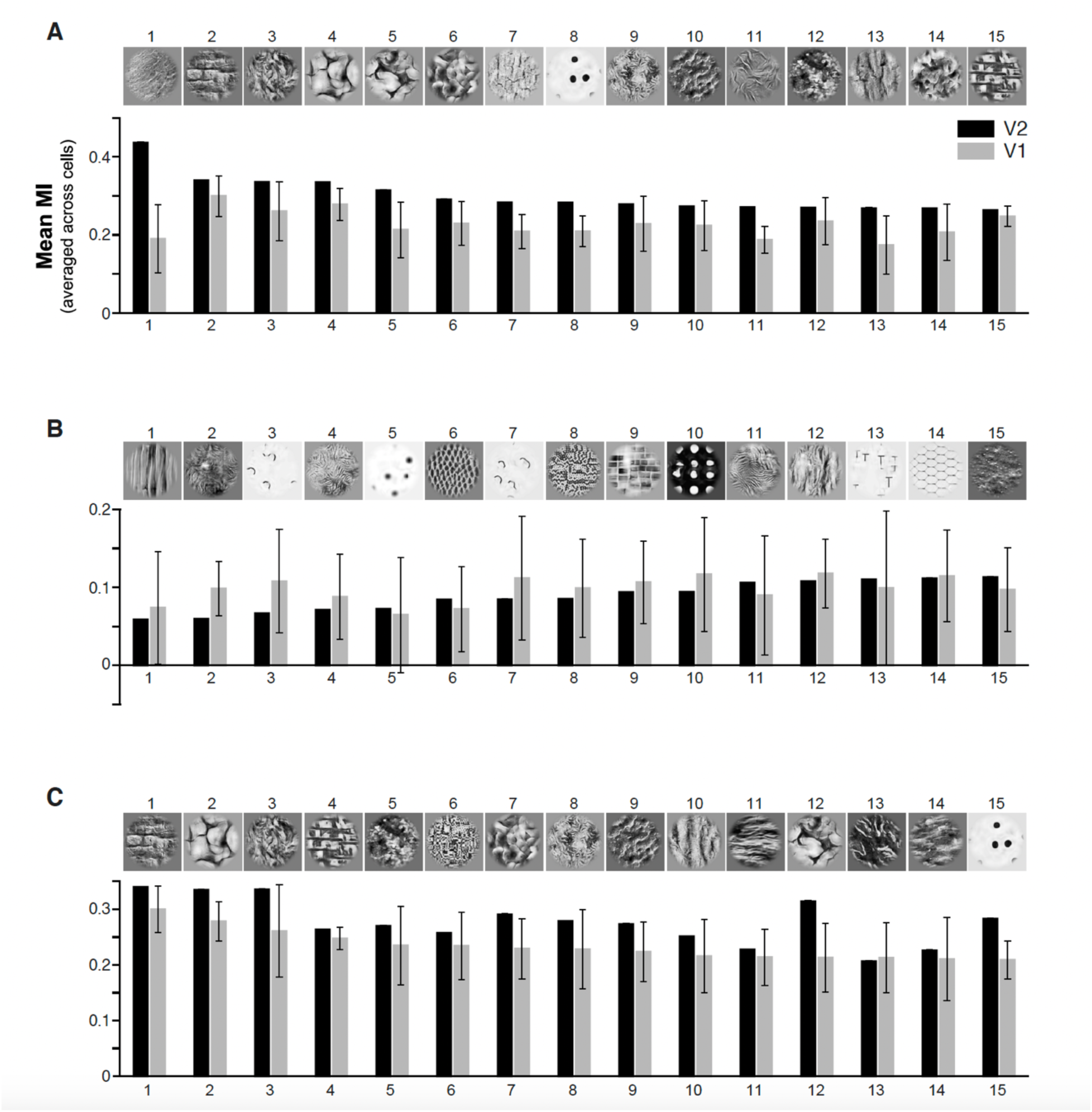
Naturalistic textures preferred by V2 and V1 cells. **(A)** Example images of the 15 naturalistic texture families most preferred by V2 cells are shown above the mean MIs (averaged across all cells) for that texture for all modeled V2 cells (n=844, *black*) and for their V1 input cells (n=11497, *gray*). Error bars: s.e.m. **(B)** Example images of the 15 naturalistic texture families least preferred by V2 cells are shown above the mean MIs for that texture for all modeled V2 and V1 cells. **(C)** Same as (A) but for the 15 naturalistic texture families most preferred by V1 cells.

## DISCUSSION

We used optical imaging of orientation and retinotopic maps in macaque visual areas V1 and V2 combined with injections of retrograde tracers in V2 orientation columns to study the functional organization of V1 inputs to V2. Single tracer injections involving one or two V2 orientation columns labeled between 162 and 7402 V1 neurons in L2/3. The aggregate RF of the labeled V1 inputs to a single injected V2 site was about the size of a single V2 RF in visual space. The V1 cells sending inputs to a single V2 site had preferred orientations that were generally biased within ±22.5° of the orientation preferred at the V2 site, but encompassed a broader range of orientations, forming complex spatial patterns that included collinear and parallel edge elements, angular and curvature configurations, and textural patterns. To understand whether and how the combination by V2 cells of information from these oriented V1 RFs could generate the more complex RF properties of V2 cells, we built a simple linear feedforward model. In this model, V2 RFs were calculated as the spatial weighted sum of their anatomically-identified V1 inputs. The resulting V2 RFs fell in two broad classes: elongated RFs with distinct ON and OFF oriented regions, resembling, but more elongated than, typical V1 cells, and complex RFs containing multiple orientations as well as non-oriented regions. Remarkably, and despite the simplifying assumptions necessitated by the limitations of our data, this simple feedforward model accurately predicted the responses of V2 orientation columns to oriented gratings, and explained the greater selectivity for naturalistic visual textures of V2 cells compared to V1 cells. Our results demonstrate that a simple linear combination of identified V1 feedforward inputs can account for the orientation-tuning of V2 RFs and their enhanced selectivity to naturalistic textures, a signature feature of area V2^32^.

Several prior studies of V2 responses have indicated that V2 RFs consist of two or more populations^30, 36–39^. We found that the distribution of POs of V1 afferents was sometimes narrow and strongly biased towards the PO of the target V2 site, particularly for the V1 inputs to V2 pale stripes, and other times was broader, representing a wider diversity of orientations, particularly so for V1 inputs to thick stripes. Consistent with this orientation-organization of the V1 afferents, our model V2 RFs generated by a linear weighted combination of these V1 afferents fell into two broad RF classes, analogous to the “uniform’ and “non-uniform” RFs previously described by Anzai et al. ^38^, or the “ultralong-Gabor” and “complex-shaped” neurons previously described by Liu et al.^30^. Consistent with our finding of narrower distributions of POs for V1 inputs to the pale stripes, Liu et al.^30^ found that V2 neurons with “ultralong-Gabor” RFs were preferentially localized within pale stripes. These authors found neurons with complex-shaped RFs to dominate in thin stripes, but our study did not investigate V1 inputs to thin stripes. Liu et al.^30^ further demonstrated that the responses to natural images of both the elongated and complex V2 RFs can be well fitted by the summation of multiple Gabor functions, and suggested a simple feedforward convergent model as the basis for the generation of V2 RFs. However, these authors lacked the anatomical data on V1-to-V2 connections that we have in our study. Thus, our study demonstrates for the first time that V2 RFs similar to those previously reported by these and other authors can be generated by a simple feedforward convergence of their anatomically-identified V1 inputs.

Any V1 input can potentially contact excitatory and/or inhibitory V2 neurons, and therefore excite or suppress V2 cells. In our model, by design, V1 inputs with positive weights were those with POs similar to the target V2 PO, while those with the negative weights were those with POs orthogonal to the V2 site’s PO. We observed a complex distribution of these positive and negative inputs with a noticeable pattern of clusters of positive iso-oriented weights flanked by clusters of negative cross-oriented weights (Fig. 4B). Previous studies based on a statistical analysis of neural responses to natural images found similar patterns of On and Off subregions within V2 RFs ^22, 26^. Such an organization may serve to enhance the local representation of excitatory signals through cross-orientation suppression.

Our linear feedforward model could accurately describe the orientation-tuning properties of V2 cells. We explored a simple-cell and a complex-cell model and found increased predictive performance using the complex cell model, more noticeably in the prediction of the width of the orientation tuning curves. This result supports Rowekamp and Sharpee’s suggestion^26^ that when phase-sensitive V1 neurons project to V2 they form quadrature pairs by projecting together with other V1 neurons tuned to other spatial phases with similar orientation/position.

A signature feature of primate area V2, that distinguishes it from V1, is its greater sensitivity to naturalistic textures^13^. Most V2 cells respond more vigorously to naturalistic textures, which contain higher-order statistical dependencies found in natural textures, than to their spectrally-matched noise images, which lack these dependencies. Consistent with these previous electrophysiological recording studies of V2, our model V2 cells on average responded more robustly to naturalistic textures compared to V1 cells, demonstrating that this complex property of V2 RFs can be generated by a linear weighted combination of feedforward V1 inputs. That feedforward mechanisms can generate V2 neuron sensitivity to naturalistic textures is also supported by experimental evidence^13^ that the greater response to textures vs. noise, measured as a modulation index (MI), is present from the onset of V2 cell responses to naturalistic stimuli. This differential response to texture vs. noise instead appears in the later part of V1 neuron responses, suggesting it may be inherited from V2 via feedback mechanisms.

Freeman et al.^13^ found lower MI values in V1 and, to a lesser extent, in V2 than we found for our model cells. This could have resulted from a number of methodological differences between the two studies. First, we tested a much wider range of texture families (n=97) than Freeman et al. (n=15). As in both studies subsets of texture families evoked greater responses from V1 and V2 cells, sampling from a wider range of textures may have led us to identify a larger number of preferred textures for V1 and V2 cells, with resulting larger MI values. Second, unlike Freeman et al., we rotated our texture images and analyzed responses to the rotation that evoked the strongest modulation. Finally, unlike Freeman et al., we do not have negative MI values, as in the complex cell model V1 and V2 responses were phase invariant. The latter two differences among the two studies likely contributed to the larger MI values in our study.

In our study, while our sample of modeled V1 and V2 cells (11,479 and 844, respectively) was substantial, the number of independent V2 cells was limited by the number of injection cases. This limitation underscores the necessity for caution in generalizing our findings, as the sampled V2 cells may not fully represent the diversity of the V2 cell population. Moreover, we made the assumption that all V1 input cells labeled by a single injection contributed to the response of each pixel at the V2 injected site. This assumption was justified by the large overlap in RF tuning properties and retinotopic location of neurons at the V2 injected column, and the observation that the aggregate RF of the entire V1 labeled field matched the average RF size of V2 neurons. However, our methods, cannot rule out that only a subset of the large labeled V1 neuron population contributed to each V2 cell RF within the injected column, or that the orientation selectivity of these subsets could provide better fits to the V2 orientation tuning.

In summary, our results demonstrate that a weighted linear combination of V1 feedforward inputs can substantially account for the more complex RF structure of V2 cells and their tuning to oriented contours and naturalistic textures.

## METHODS

### Animals

All procedures were approved by the University of Utah’s Animal Care and Use Committee and were in accordance with National Institute of Health and USDA guidelines. Four adult macaque monkeys (*Macaca fascicularis*; 2 males, 2 females; 3-6.5 kg) were used in this study. Animals were purchased from a commercial breeder, quarantined for 6 weeks and group-housed at the University of Utah prior to being used for experimentation. In 3 of the 4 animals, optical imaging and tracer injections were performed during a single 4 day-long anesthetic event, at the end of which the animal was euthanized. In the fourth animal (MK368), a quick (2-3hrs long) imaging session was performed under anesthesia to obtain functional maps and identify the V2 CO stripes, the tracers were then injected, and the animal recovered from anesthesia. Four days later this animal was re-anesthetized and maintained under anesthesia for an additional 4 days during which additional optical imaging was performed and the animal was euthanized at the end of the 4^th^ day.

### Surgical procedures

Animals were pre-anesthetized with Ketamine (10-20 mg/kg i.m.), intubated with an endotracheal tube and artificially ventilated. During surgery, anesthesia was maintained with isoflurane (0.5–2.5%) in 100% oxygen. End-tidal CO_2_, blood O_2_ saturation, electrocardiogram, blood pressure, lung pressure, and body temperature were monitored continuously. The animal’s head was fixed in a stereotaxic apparatus. The scalp was incised, a large craniotomy and durotomy were made to expose the lunate sulcus and areas V2 and V1 posterior to it. A clear sterile silicone artificial dura was placed on the cortex, and the craniotomy was filled with a sterile 3% agar solution and sealed with a sterile glass coverslip glued to the skull with Glutures (Abbott Laboratories, Lake Bluff, IL). On completion of the surgery, the isoflurane was turned off and anesthesia was maintained with sufentanil citrate (5-10 µg/kg/h, i.v.). The pupils were dilated with a long-acting topical mydriatic agent (atropine; 3 animals) or a short-acting one (tropicamide; in 1 animal that was recovered after surgery), the corneas protected with gas-permeable contact lenses, the eyes refracted, and optical imaging was started. In one animal (MK368), optical imaging was performed for about 2 hours to obtain orientation maps and identify the V2 stripes. Then the glass coverslip, agar and artificial dura were removed, and the tracer were injected into V2, targeted at specific orientation domains and V2 stripe types using the surface vasculature as guidance. We only made injections in V2 thick or pale stripes, as, unlike thin stripes, these contain well defined orientation maps. On completion of the injections, new artificial dura was placed on the cortex, the craniotomy was filled with Gelfoam and sealed with sterile parafilm and dental cement, the skin was sutured, and the animal was recovered from anesthesia. After a 4 day survival period, this animal was again anesthetized, a new optical window was made over the craniotomy as described above and imaging of V1 and V2 was performed continuously for 4 days under sufentanil anesthesia and paralysis (vecuronium bromide, 0.1-0.3µg/kg/h, i.v.) to stabilize the eyes, at the end of which the animal was euthanized with Beuthanasia (0.22 ml/kg, i.v.) and perfused transcardially with saline for 2–3 min, followed by 4% paraformaldehyde in 0.1M phosphate buffer for 20-25 min (thus post-injection survival time for this animal was 8 days).

In the other 3 animals, initial optical imaging of V1 and V2 was performed for up to 12 hours, the chamber was then removed, and the tracer injections made in V2 thick and pale stripes. After the injections a new optical window was made and imaging was continued under anesthesia for an additional 3-3.5 days post-injections, after which they animals were euthanized and perfused as described above.

### Tracer injections

A total of 12 retrograde tracer injections were made in 4 animals (2-4 injections per animal in the same hemisphere). We excluded 2 injection sites from analysis because they spanned too many orientation columns for the purposes of our study. The tracers consisted of Cholera Toxin B (CTB)-alexa conjugated to different fluorophores (647, 488, 555). In animals that received 4 injections in the same hemisphere, two injections of the same tracer (CTB-555) were spaced by at least 5 mm to ensure no overlap of the resulting labeled fields in V1. The tracers were pressure injected using picospritzer and a glass micropipette (30-40 µm tip). The pipette was lowered to a depth of 600-800 µm from pia, with the goal of targeting layers 3-4 (where the majority of V1 inputs terminate in V2) and 30-45nl of tracer solution (3% in distilled water) were slowly injected. The pipette was left in place for an additional 10 minutes before being retracted.

### Intrinsic signal Optical imaging

We performed intrinsic signal optical imaging (OI) using the Imager 3001 and VDAQ software (Optical Imaging Ltd, Israel^40^) under red light illumination (630 nm). The surface vasculature was captured using green light (546 nm) illumination (as in **Extended Data** Fig. 1A) and used as guidance to target injections to specific functional domains *in vivo*, as well as for *ex vivo* co-registration of the functional images with the histological sections (**Extended Data** Fig. 1). Data acquisition rate was 5Hz.

### Visual Stimuli

Orientation maps were obtained by presenting binocularly full-field, high-contrast (100%), pseudorandomized achromatic drifting square-wave gratings of 8 orientations (0°–horizontal, 22.5°, 45°, etc.), spatial frequency of 0.5-2.0 cycles/° and temporal frequency of 1.5 cycles/s, drifting back and forth, orthogonal to the grating orientation. Each grating orientation was presented 30 times, resulting in a total of 240 trials. Retinotopic maps were obtained by presenting monocularly oriented gratings (horizontal or vertical) occupying complementary adjacent strips of visual space, i.e. masked by 0.5-1° strips of uniform gray repeating every 0.5-1°, with the masks reversing in position in alternate trials (Fig. 1 **H,I**), for a total of 60 trials. Each trial lasted 6s, and consisted of 1s of gray screen, 4s of stimulus presentation, 1s of stimulus off.

### Histology

After perfusion the block containing areas V1 and V2 was post-fixed between glass slides for 1-2 hrs. to slightly flatten the cortex in the imaged area, then sunk in 30% sucrose for cryoprotection, and frozen-sectioned at 40 μm tangentially, parallel to the plane of optical imaging. Sections were mounted and coverslipped and imaged for fluorescent label using a confocal microscope. Every third section was reacted free-floating for cytochrome oxidase (CO) staining and imaged on a Zeiss Axio Imager Z2 light microscope. Imaged CO sections were aligned to images of fluorescent label in adjacent sections using blood vessels as guidance. Histological sections were aligned to *in vivo* imaged functional maps as described in the *Data Analysis* section below.

### Data analysis

#### Analysis of optical imaging data

##### Retinotopic Maps

Responses to visual stimuli were averaged frame by frame across all trials resulting in a single stack for each stimulus condition. Frames were binned into 1-second frames and the first frame was subtracted from the rest to remove baseline signal. Difference images were obtained by subtracting two complementary conditions (reversed masks stimuli). Extended spatial decorrelation (ESD)^41^ analysis was applied to difference stacks in order to separate stimulus evoked changes in signal from biological noise and other imaging artifacts. The ESD component that showed the relevant response was used as the retinotopic map. Retinotopic stripes were hand traced (Fig. 1H**-I**) by three different individuals.

##### Computation of orientation tuning curves from optical imaging data

For each trial, a response map was measured by subtracting the baseline map (average of first two frames) from the stimulus-evoked map (average of frames 15 to 20). These response maps were then inverted (multiplied by -1), so that the bright patches corresponded to the domains most responsive to the stimulus. Pixel intensity values were cut between +/-100 (pixels values above/below +/-100 were set to +/-100, respectively). The resulting maps were divided by a control image, consisting of the average of the first five frames of the first recorded trial. To reduce low spatial frequency noise, these maps were, then, high-pass filtered by subtracting low frequency noise maps (maps smoothed by an isotropic 2D Gaussian kernel with standard deviation equal to 25 pixels ≈ 0.45 *mm*) from smoothed response maps (smoothed by Gaussian kernel with standard deviation equal to 2.5 pixels ≈ 45 *µm*). Responses to each grating orientation were calculated by averaging across trials for that orientation, excluding the first trial. A stimulus non-specific response map (minimum response map across eight orientation maps) was subtracted from each stimulus response map. Finally, these single condition maps were rescaled so that pixel values in the map ranged between 0 and 100. This was done by dividing each map by the maximum of the difference map (calculated by subtracting the minimum map - across orientations - from the maximum map, and excluding blood vessels) and multiplied by 100 following established procedures^42^. Tuning curves were finally estimated by fitting Von Mises function to the data using the Levenburg–Marquardt algorithm^43, 44^. Functional maps were up-sampled by 20x to match the size of the microscopy images of tissue sections. Blood vessels were segmented semi-automatically using a script kindly provided by Dr. Amir Shmuel (Brain Imaging Signals Lab, McConnell Brain Imaging Centre, Montreal Neurological Institute).

##### Orientation Preference Maps

Orientation preference maps were generated through pixel-by-pixel vector summation of the responses to eight grating orientations (8 single condition maps). The PO of each pixel was the angle of the resultant vector^45^.

##### Identification of V2 stripes

To target injections of tracers to specific V2 stripes, the latter were identified on the functional maps imaged *in vivo* as follows. Thick stripes were identified as the middle of regions having an orientation-preference map, and pale stripes as regions having an orientation-preference map immediately neighboring a striped region with weak or no systematic orientation maps (the latter corresponding to thin stripes (**Extended Data** Fig. 1E). Stripe identity and borders were then confirmed postmortem by aligning CO-stained sections to the functional maps as described below. On CO-stained tissue sections, dark CO stripes were classified as thick or thin if they appeared thicker or thinner, respectively, than the two adjacent dark stripes and coincided with thick or thin stripes, respectively, as defined in the optical imaging maps (**Extended Data** Fig. 1D, E).

#### Analysis of anatomical data

##### Alignment of histological sections to in vivo functional maps

The retinotopic position and orientation preference of V1 labeled cells and V2 injection sites were based on these cell/pixel locations on the functional maps. To this purpose, functional maps and histological tissue sections were aligned using the surface vasculature as guidance. Specifically, the most superficial tangential tissue section containing the surface vessels (which run parallel to the cortical surface; **Extended Data** Fig. 1B) was warped to the image of the cortical surface vasculature obtained *in vivo* under green light (the latter in register with the functional maps obtained under red light; **Extended Data** Fig. 1A,E). **Extended Data** Fig. 1C shows the two warped images co-registered. Each deeper tissue section was then registered sequentially to the top sections by aligning the radial blood vessels (which run perpendicular to the cortical surface). This alignment procedure is illustrated in **Extended Data** Fig. 1. Warping of tissue sections to functional maps was performed using IR-tweak warping software (NCR toolset, Scientific Computing and Imaging Institute, University of Utah). IR-tweak is an interactive, multithreaded, application for manual slice-to-slice registration. As control points are placed by the user in one image, their locations in the other image are estimated by the current thin-plate spline transform parameters^46^.

##### Plotting labeled cells and injection sites

An experienced observer marked by hand a single pixel at each V1 labeled cell location in the stack of aligned tissue sections through L2/3, and the marked cells were overlaid to the functional maps (e.g. Fig. 1C**,H**). Tracer injection sites in V2 were outlined on multiple images of injection sites through the depth of the cortex; a composite injection site encompassing all outlines in individual sections was overlaid onto the merged stack, and its size and V2 stripe location were determined. The size of the CTB-alexa injection site (the tracer uptake zone) was defined as the dense core seen under fluorescence microscopy, within which no labeled cell bodies were discernible^47^ (e.g. Fig. 1F).

#### Cortex to visual field mapping

In order to locate the RF locations of the labeled V1 cells in the visual field, we used retinotopic maps co-registered to the histological sections. An area encompassing all V1 labeled cells with borders drawn parallel to the imaged horizontal and vertical retinotopic stripes was defined (yellow contour in Fig. 1H**,I**). The V1-V2 border (extracted from co-registered CO images), corresponding to the representation of the vertical meridian, served as one of four sides of this area. The size of the area was estimated in degrees of visual angle based on the number and size of the retinotopy stripes it encompassed, which corresponded to the known width of the stripes in the visual stimulus used to activate the cortex (insets in Fig. 1H**,I**). We used an Elliptic grid generation^29^ approach for dividing this odd-shaped area into a grid and registering it to a grid in visual space (Fig. 1 **J,K**). In this approach, in an iterative way (2500000 iterations and convergence error <10^-5^), the space is divided into smaller pieces until an evenly distributed grid is achieved (more details are provided in **Supplementary Methods**). Each cell’s retinotopic location in visual space was estimated from its closest proximity to the grid points in the cortex (Fig. 1K).

#### Statistics

We performed several statistical tests to examine whether the observed distributions of preferred orientation (PO) of the labeled V1 cells were significantly different from distributions simulated to test particular selectivity hypotheses. As described in the Results and in **Extended Data** Fig. 4, we performed 3 kinds of statistical comparisons. First, we compared the observed distributions of the V1 labeled cells’ POs with the distribution of POs of all the pixels within the V1 labeled field (**Extended Data** Fig. 4A, B **top**), and of all the pixels within the V1 imaged area (Fig. 1E **and Extended Data** Fig. 4F **top**), using a chi-square goodness of fit test. We also compared the observed distribution of the V1 labeled cells’ POs with simulated distributions that we obtained by two different random placement strategies. In the first test, control distributions were simulated by resampling M pixels (M = number of labeled V1 cells) 1,000 times within the labeled field area (**Extended Data** Fig. 4A). Summary statistics were calculated for observed and simulated distributions. Because of the circular nature of the orientation data, circular statistics including mean resultant length (MRL) associated with the mean orientation and circular standard deviation (CSD) - as described by Fisher (1993)^31^ - were used. MRL and CSD from observed distributions were compared to distributions of those statistics from simulated distributions. Furthermore, observed and simulated distributions were compared using the Kolmogorov–Smirnov test at a Bonferroni-corrected family-wise p-value of 0.05 (**Extended Data** Fig. 4D). In the second test, control distributions were obtained by shifting randomly (> 1500 times and as much as the field of view/ imaging area allowed) the observed pattern of cell label across the V1 orientation map for that case (**Extended Data** Fig. 4E,H). Circular statistics (MRL and CSD; **Extended Data** Fig. 4I) and the Kolmogorov–Smirnov test were calculated as described above.

To understand how the orientation organization of V1 to V2 connections differs from a perfect like-to-like connectivity (i.e. V1 cells connecting exclusively with V2 cells having the same PO), we simulated these distributions under like-to-like connectivity and compared them to the observed distributions **(Extended Data** Fig. 5). We postulated that under like-to-like connectivity, if the retrograde tracer injection sites were precisely confined to a single orientation column, the resulting distribution of POs of the labeled V1 cells would resemble a Gaussian function. This is because POs are derived from orientation maps that undergo spatial smoothing with a Gaussian kernel (as described above). We used 22.5° for full width at half maximum because the orientation responses were experimentally sampled at a resolution of 22.5°. In most experiments, however, the tracer injections were not perfectly confined to a single V2 orientation column, therefore the join probability distribution of V1 cells POs (*p*(*θ*_1_, …*θ_N_*\*)) were simulated as a weighted sum of Gaussian functions centered at the PO of the injection site as described by (Eq.1) and illustrated in **Extended Data** Fig. 5A:

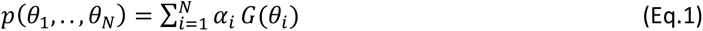

Where *G*(*θ*_0_) is Gaussian function centered at *θ*_0_ and *α*_0_ is the fraction of pixels at the injection site that have PO of *θ*_0_. *N* is the number of orientation columns involved by the injection site. The simulated distributions were binned similarly to the observed data, and we conducted a Chi-square analysis of the binned data, to determine the similarity of the two distributions. If the test didn’t refute the null hypothesis, it indicated that V1-to-V2 connections follow a perfect like-to-like connectivity. Our results indicated otherwise.

### Modeling V2 cells responses to visual stimuli

The responses of V2 cells to visual stimuli were modeled as a linear weighted sum of the responses of their V1 inputs. Given the absence of data on the phase sensitivity of V1 cells in our imaging, V1 cells were modeled (i) as phase-sensitive simple cells all with either even-parity RFs or odd-parity RFs, or (ii) as phase-invariant complex cells. Two dimensional (2D) Gabor functions (Eq.2) were used to model simple cells’ RFs^48, 49^, and their response to an image (visual stimulus) was estimated by half-wave rectifying the inner dot product of the image and the RF. The estimated retinotopic location of each cell in visual space (determined as described above) was used for the center coordinates of the Gabor function (*x_c_, y_c_*), and the cell’s PO was used for orienting the Gabor function (*θ*_4_). To estimate the aspect ratio of the cell’s RF (*γ*_4_), we first simulated the responses of a set of Gabor functions with 40 different aspect ratios, ranging from 0.1 to 4, to grating stimuli of 8 different orientations, and measured their orientation tuning curves. Next, these tuning curves were fit to each V1 cell’s tuning curve computed from our OI data (see above for details), and the aspect ratio was selected based on the goodness of fit. The standard deviation of the elliptical Gaussian along *x* (σ_x_) was set to ≈ 0.6 corresponding to an optimal spatial frequency (*s_f_*) of 1 cycles per degree and a bandwidth of 1 octave^50^.

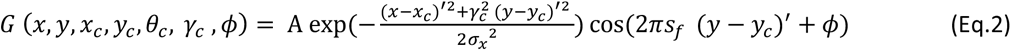

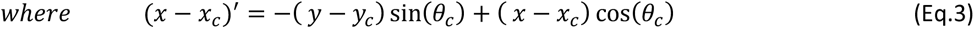

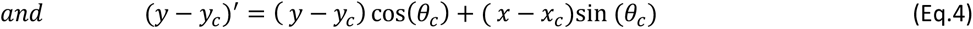

In equation 2 above, A is a normalization factor that sets the L2 norm of the Gabor function to 1, and *ϕ* is 0 for even-parity and π/2for odd-parity RFs.

The response of a V1 complex cell to an image was measured by summing half-wave rectified responses of four simple cells with RFs modeled as Gabor filters with identical parameters except for spatial phase that was offset by 90 degree (Fig. 4A)^33^.

Weights in the linear model were estimated for each V1-V2 cell/pixel pair as the dot product of their mean-subtracted and normalized tuning curves. Specifically, this was estimated by considering each tuning curve as an 8-dimensional vector, subtracting the mean, normalizing by dividing to the vector’s length, and calculating the two vectors inner products:

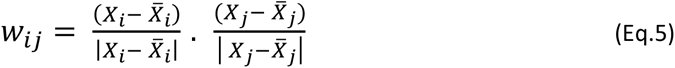

where, *w_ij_* is the weight for *i*th V1 cell and *j*th V2 pixel pair, *X_i_* & *X_j_* and *X̄_i_* & *X̄_i_* are 8 dimensional vectors representing the tuning curve and its mean for given V1 cell and V2 pixel respectively. |*X*| indicates the magnitude of the vector *X*. Subtracting the mean and normalizing the tuning curves results in negative weights for the pairs with orthogonal PO (maximum weight = -1) and positive weight (maximum weight = +1) for those with similar orientations. The motivation for using this method for measuring weighs was to replicate the cross-orientation organization of local edges previously described for V2 RFs^26^ (see Results and Discussion).

Model performance was tested by eight-fold leave-one-out cross validation process ^51^. Model error was quantified as mean relative error (with respect to the measured response range).

#### Texture images

Texture images were synthesized from 228 texture photographs obtained from the online collections by Phil Brodatz (https://www.ux.uis.no/~tranden/brodatz.html), Javier Portilla and Eero P. Simoncelli (http://www.cns.nyu.edu/~lcv/texture/index.php), and VisTex, MIT Media Laboratory (https://vismod.media.mit.edu/vismod/imagery/VisionTexture/vistex.html) using a software package (https://github.com/LabForComputationalVision/textureSynth) written by Javier Portilla and Eero Simoncelli^35, 52, 53^. These 228 original images were converted to grayscale and resized to have 256x256 pixels, and texture parameters were extracted using a ‘textureAnalysis’ code that processed the images with a multi-scale, multi-oriented bank of filters with 4 orientations, 4 spatial scales, and in a 9 by 9 local neighborhood. Naturalistic textures were synthesized using the ‘textureSynthesis’ code with the number of iterations set to 25, and output image size set to 192x128 pixels. These synthesized images were then cropped to a square shape and resized to 320x320 pixels, and a circular mask (diameter = 320 pixel) was applied (in our analysis 100 pixels correspond to 1° in visual field). For each original image, 30 naturalistic texture images (NTI) and 30 corresponding spectrally-matched noise images (SMNI) were synthesized^13^. Each SMNI was generated by randomizing the phase values of the Fourier transform of the original image and then inverting the Fourier transforms. A user was asked to look at the 30 samples and rate the “naturalness” of the synthesized textures from 1 to 10; the textures with a naturalness score <8 were discarded. From the 228 synthesized texture families, 97 passed the test. These images were rotated 8 times around the clock in 22.5° steps, resulting in 240 texture and 240 noise samples within each texture family.

### Analysis of responses to textures in the model

V1 and V2 RFs and images were standardized to have zero mean and unit standard deviation before calculating their dot product. For each texture and noise pair, a modulation index (MI) was defined as the response to the texture minus the response to the noise divided by their sum. To compare across different cells, we opted to use results at the image rotation that provided the most significant differential response (i.e. the smallest relative variance, in other words, the largest across sample mean over variance value). This approach ensures that when comparing across different cells, the effect of oriented features in the responses are similarly considered and the comparison reflects the effect of higher-order statistical dependencies that are common across different samples of a naturalistic texture.

## DATA AVAILABILITY STATEMENT

The data presented here will be provided upon request to the corresponding authors. Source data for the figures are provided with the paper.

## Supporting information

Supplementary Figurers

## ACKNOWLEDGMENTS

We thank Kesi Sainsbury for histological assistance. Dr Amir Shmuel kindly provided us with a MATLAB code for semiautomatic segmentation of blood vessels and provided advice on the processing of optical images. The MATLAB code for analyzing and synthesizing visual textures was written by Javier Portilla and Eero Simoncelli, 1999-2000 (https://github.com/LabForComputationalVision/textureSynth/tree/master). This work was supported by grants from the National Institute of Health (NIH) to A.A. (R01 EY026812, R01 EY019743, R01 EY031959), and to Q.Z. (R21 EY035085) the National Science Foundation (IOS 1755431) and the Mary Boesche Endowed Chair, to A.A; an unrestricted grant from Research to Prevent Blindness, Inc., New York, NY, a Core grant from the NIH (P30 EY014800), and a Training grant from the NIH (T32 EY024234) to the Department of Ophthalmology & Visual Sciences, University of Utah. The departmental Training grant was awarded to M.S.H..

## AUTHOR CONTRIBUTIONS

Conceptualization: M.S.H., S.M., Q.Z.,A.A. Investigation: S.M., F.F., A.A. Data Analysis: M.S.H., S.M. Modeling: M.S.H., Q.Z. Writing-Original Draft: M.S.H., A.A., Q.Z. Writing–Review/Editing: M.S.H., A.A., Q.Z. Visualization: M.S.H., A.A. Supervision & Funding Acquisition: A.A., Q.Z.

## REFERENCES

1. Van Essen, D.C. & Maunsell, J.H.R. Hierarchical organization and functional streams in the visual cortex. Trends Neurosci. 6, 370–375 (1983). doi:10.1016/0166-2236(83)90167-4

2. Blasdel, G.G. Orientation selectivity, preference, and continuity in monkey striate cortex. J Neurosci 12, 3139–3161 (1992). doi:10.1523/JNEUROSCI.12-08-03139.1992

3. Hubel, D.H. & Wiesel, T.N. Receptive fields and functional architecture of monkey striate cortex. J Physiol 195, 215–243 (1968). doi:10.1113/jphysiol.1968.sp008455

4. Desimone, R., Albright, T.D., Gross, C.G. & Bruce, C. Stimulus-selective properties of inferior temporal neurons in the macaque. J Neurosci 4, 2051–2062 (1984). doi:10.1523/JNEUROSCI.04-08-02051.1984

5. Tanaka, K., Saito, H., Fukada, Y. & Moriya, M. Coding visual images of objects in the inferotemporal cortex of the macaque monkey. J Neurophysiol 66, 170–189 (1991). doi:10.1152/jn.1991.66.1.170

6. Freiwald, W.A. & Tsao, D.Y. Functional compartmentalization and viewpoint generalization within the macaque face-processing system. Science 330, 845–851 (2010). doi:10.1126/science.1194908

7. Markov, N.T., et al. A weighted and directed interareal connectivity matrix for macaque cerebral cortex. Cereb Cortex 24, 17–36 (2014). doi:10.1093/cercor/bhs270

8. Van Essen, D.C., Newsome, W.T., Maunsell, J.H. & Bixby, J.L. The projections from striate cortex (V1) to areas V2 and V3 in the macaque monkey: asymmetries, areal boundaries, and patchy connections. J Comp Neurol 244, 451–480 (1986). doi:10.1002/cne.902440405

9. Vanni, S., Hokkanen, H., Werner, F. & Angelucci, A. Anatomy and Physiology of Macaque Visual Cortical Areas V1, V2, and V5/MT: Bases for Biologically Realistic Models. Cereb Cortex 30, 3483–3517 (2020). doi:10.1093/cercor/bhz322

10. Girard, P. & Bullier, J. Visual activity in area V2 during reversible inactivation of area 17 in the macaque monkey. J Neurophysiol 62, 1287–1302 (1989). doi:10.1152/jn.1989.62.6.1287

11. Hegde, J. & Van Essen, D.C. Selectivity for complex shapes in primate visual area V2. J Neurosci 20, RC61 (2000). doi:10.1523/JNEUROSCI.20-05-j0001.2000

12. Ito, M. & Komatsu, H. Representation of angles embedded within contour stimuli in area V2 of macaque monkeys. J Neurosci 24, 3313–3324 (2004). doi:10.1523/JNEUROSCI.4364-03.2004

13. Freeman, J., Ziemba, C.M., Heeger, D.J., Simoncelli, E.P. & Movshon, J.A. A functional and perceptual signature of the second visual area in primates. Nat Neurosci 16, 974–981 (2013). doi:10.1038/nn.3402

14. Ziemba, C.M., Freeman, J., Movshon, J.A. & Simoncelli, E.P. Selectivity and tolerance for visual texture in macaque V2. Proc Natl Acad Sci U S A 113, E3140–3149 (2016).

15. Zhou, H., Friedman, H.S. & von der Heydt, R. Coding of border ownership in monkey visual cortex. J Neurosci 20, 6594–6611 (2000). doi:10.1523/JNEUROSCI.20-17-06594.2000

16. von der Heydt, R., Zhou, H. & Friedman, H.S. Representation of stereoscopic edges in monkey visual cortex. Vision Res 40, 1955–1967 (2000). doi:10.1016/S0042-6989(00)00044-4

17. Levitt, J.B., Kiper, D.C. & Movshon, J.A. Receptive fields and functional architecture of macaque V2. J Neurophysiol 71, 2517–2542 (1994). doi:10.1152/jn.1994.71.6.2517

18. Smith, A.T., Singh, K.D., Williams, A.L. & Greenlee, M.W. Estimating receptive field size from fMRI data in human striate and extrastriate visual cortex. Cereb Cortex 11, 1182–1190 (2001). doi:10.1093/cercor/11.12.1182

19. DiCarlo, J.J., Zoccolan, D. & Rust, N.C. How does the brain solve visual object recognition? Neuron 73, 415–434 (2012). doi:10.1016/j.neuron.2012.01.010

20. Marr, D. Vision : a computational investigation into the human representation and processing of visual information (W.H. Freeman, San Francisco, 1982).

21. Pasupathy, A. & Connor, C.E. Shape representation in area V4: position-specific tuning for boundary conformation. J Neurophysiol 86, 2505–2519 (2001). doi:10.1152/jn.2001.86.5.2505

22. Hosoya, H. & Hyvarinen, A. A Hierarchical Statistical Model of Natural Images Explains Tuning Properties in V2. J Neurosci 35, 10412–10428 (2015). doi:10.1523/JNEUROSCI.5152-14.2015

23. Hoyer, P.O. & Hyvarinen, A. A multi-layer sparse coding network learns contour coding from natural images. Vision Res 42, 1593–1605 (2002). doi:10.1016/S0042-6989(02)00017-2

24. Hyvarinen, A., Gutmann, M. & Hoyer, P.O. Statistical model of natural stimuli predicts edge-like pooling of spatial frequency channels in V2. BMC Neurosci 6, 12 (2005). doi:10.1186/1471-2202-6-12

25. Karklin, Y. & Lewicki, M.S. Learning higher-order structures in natural images. Network 14, 483–499 (2003). doi:10.1088/0954-898X_14_3_306

26. Rowekamp, R.J. & Sharpee, T.O. Cross-orientation suppression in visual area V2. Nat Commun 8, 15739 (2017). doi:10.1038/ncomms15739

27. Ts’o, D.Y., Roe, A.W. & Gilbert, C.D. A hierarchy of the functional organization for color, form and disparity in primate visual area V2. Vision Res. 41, 1333–1349 (2001). doi:10.1016/S0042-6989(01)00076-1

28. Felleman, D.J., et al. The Representation of Orientation in Macaque V2: Four Stripes Not Three. Cereb. Cortex 25, 2354–2369 (2015). doi:10.1093/cercor/bhu033

29. Spekreijse, S.P. Elliptic Grid Generation Based on Laplace Equations and Algebraic Transformations. Journal of Computational Physics 118, 38–61 (1995). doi:10.1006/jcph.1995.1078

30. Liu, L., et al. Spatial structure of neuronal receptive field in awake monkey secondary visual cortex (V2). Proc Natl Acad Sci U S A 113, 1913–1918 (2016). doi:10.1073/pnas.1525505113

31. Fisher, N.I. Statistical Analysis of Circular Data. Cambridge University Press, London. (1993).

32. Freeman, J., Ziemba, C.M., Heeger, D.J., Simoncelli, E.P. & Movshon, J.A. A functional and perceptual signature of the second visual area in primates. Nat. Neurosci. 16, 974–981 (2013). doi:10.1038/nn.3402

33. Lian, Y., et al. Learning receptive field properties of complex cells in V1. PLoS Comput Biol 17, e1007957 (2021). doi:10.1371/journal.pcbi.1007957

34. Movshon, J.A., Thompson, I.D. & Tolhurst, D.J. Spatial summation in the receptive fields of simple cells in the cat’s striate cortex. J Physiol 283, 53–77 (1978). doi:10.1113/jphysiol.1978.sp012488

35. Portilla, J. & Simoncelli, E.P. A parametric texture model based on joint statistics of complex wavelet coefficients. International journal of computer vision 40, 49–70 (2000). doi:10.1023/A:1026553619983

36. Schmid, A.M., Purpura, K.P., Ohiorhenuan, I.E., Mechler, F. & Victor, J.D. Subpopulations of neurons in visual area v2 perform differentiation and integration operations in space and time. Front Syst Neurosci 3, 15 (2009). doi:10.3389/neuro.06.015.2009

37. Schmid, A.M., Purpura, K.P. & Victor, J.D. Responses to orientation discontinuities in V1 and V2: physiological dissociations and functional implications. J Neurosci 34, 3559–3578 (2014). doi:10.1523/JNEUROSCI.2293-13.2014

38. Anzai, A., Peng, X. & Van Essen, D.C. Neurons in monkey visual area V2 encode combinations of orientations. Nat Neurosci 10, 1313–1321 (2007). doi:10.1038/nn1975

39. Willmore, B.D., Prenger, R.J. & Gallant, J.L. Neural representation of natural images in visual area V2. J Neurosci 30, 2102–2114 (2010). doi:10.1523/JNEUROSCI.4099-09.2010

40. 40. Grinvald, A., et al. In-vivo optical imaging of cortical architecture and dynamics. Modern Techniques in Neuroscience Research (1999).

41. Schiessl, I., et al. Blind signal separation from optical imaging recordings with extended spatial decorrelation. IEEE Trans Biomed Eng 47, 573–577 (2000). doi:10.1109/10.841327

42. Swindale, N.V., Grinvald, A. & Shmuel, A. The spatial pattern of response magnitude and selectivity for orientation and direction in cat visual cortex. Cereb Cortex 13, 225–238 (2003). doi:10.1093/cercor/13.3.225

43. Swindale, N.V. Orientation tuning curves: empirical description and estimation of parameters. Biol Cybern 78, 45–56 (1998). doi:10.1007/s004220050411

44. Levenberg, K. A Method for the Solution of Certain Non-Linear Problems in Least Squares. Quarterly of Applied Mathematics 2, 164–168 (1944). doi:10.1090/qam/10666

45. Bonhoeffer, T. & Grinvald, A. Iso-orientation domains in cat visual cortex are arranged in pinwheel-like patterns. Nature 353, 429–431 (1991). doi:10.1038/353429a0

46. Anderson, J.R., et al. A computational framework for ultrastructural mapping of neural circuitry. PLoS Biol 7, e1000074 (2009). doi:10.1371/journal.pbio.1000074

47. Federer, F., Williams, D., Ichida, J.M., Merlin, S. & Angelucci, A. Two projection streams from macaque V1 to the pale cytochrome oxidase stripes of V2. J. Neurosci. 33, 11530–11539 (2013). doi:10.1523/JNEUROSCI.5053-12.2013

48. Daugman, J.G. Two-dimensional spectral analysis of cortical receptive field profiles. Vision Res 20, 847–856 (1980). doi:10.1016/0042-6989(80)90065-6

49. Daugman, J.G. Uncertainty relation for resolution in space, spatial frequency, and orientation optimized by two-dimensional visual cortical filters. J Opt Soc Am A 2, 1160–1169 (1985). doi:10.1364/JOSAA.2.001160

50. Lee, T.S. Image Representation Using 2D Gabor Wavelets. IEEE TRANSACTIONS ON PATTERN ANALYSIS AND MACHINE INTELLIGENCE 18, 959–971 (1996).

51. Hastie, T., Tibshirani, R. & Friedman, J.H. *The Elements of Statistical Learning Data Mining, Inference, and Prediction* (Springer, New York, 2009).

52. Portilla, J. & Simoncelli, E.P. Texture modeling and synthesis using joint statistics of complex wavelet coefficients. in IEEE workshop on statistical and computational theories of vision (1999).

53. Portilla, J. & Simoncelli, E.P. Texture representation and synthesis using correlation of complex wavelet coefficient magnitudes (Citeseer, 1999).

