## Supplementary Methods for "Primate V2 Receptive Fields Derived from Anatomically Identified Large-Scale V1 Inputs"

**Elliptic Grid Generation.** Elliptic grid generation is one of several methods used to generate structured grids for odd geometries. This approach works well particularly for domains where all the physical boundaries are specified. In this approach a pair of Laplace equations, which is a system of elliptic equations, is solved using an iterative numerical scheme (here, Gauss-Seidel with successive over relaxation). The final solution determines the location of the interior grid point given a set of predetermined boundary points.

For this purpose, consider two separate spaces including (1) physical and (2) computational spaces. Physical space is the real-world $x - y$ space (cortex) which can be mapped to an abstract rectangular space (visual field), which we call computational space. The pair of coordinates defining a two-dimensional computational space is commonly denoted as $(\eta, \zeta)$. Here, the following pair of Laplace equation provides the aforementioned mapping from the computational to the physical domain:

$$\begin{aligned} \zeta_{xx} + \zeta_{yy} &= 0 \\ \eta_{xx} + \eta_{yy} &= 0' \end{aligned}$$

such that a uniform mesh in the computational space can be mapped to the physical space with uniformly distributed (equidistributed) nodes and grid lines which are locally perpendicular to the boundaries. In order to solve these Laplace equations, first we need to interchange the independent $(\eta, \zeta)$ and dependent variables $(x, y)$ through applying a simple mathematical transformation. Remember we know what the computational coordinates are: (1) they vary between zero and one; (2) we are the one who sets $\Delta\zeta$ and $\Delta\eta$ (or number of nodes); and they produce a uniform grid in a one-by-one square domain. Thus, in essence we are mapping a square domain to our desired odd geometry. The resultant nonlinear partial differential equation can be written as:

$$\begin{aligned} a x_{\zeta\zeta} - 2b x_{\zeta\eta} + c x_{\eta\eta} &= 0 \\ a y_{\zeta\zeta} - 2b y_{\zeta\eta} + c y_{\eta\eta} &= 0' \end{aligned}$$

Where $a$, $b$ and $c$ are functions of physical coordinates derivatives. Here, the resultant mesh adapts to the boundary of the physical space as the boundaries are fed to the solution in the form of Dirichlet boundary conditions. The simplest way to solve these equations is using an iterative method which is basically another form of the traditional trial and error scheme. First, we guess the results and next we plug them into the equations in an iterative fashion to correct our guess. Finally, the outcome of this solution is an equidistibuted node distribution with respect to the given boundary (e.g. **Fig. 1J**).
