## Supplementary Figurers for "Primate V2 Receptive Fields Derived from Anatomically Identified Large-Scale V1 Inputs"

### EXTENDED DATA FIGURES

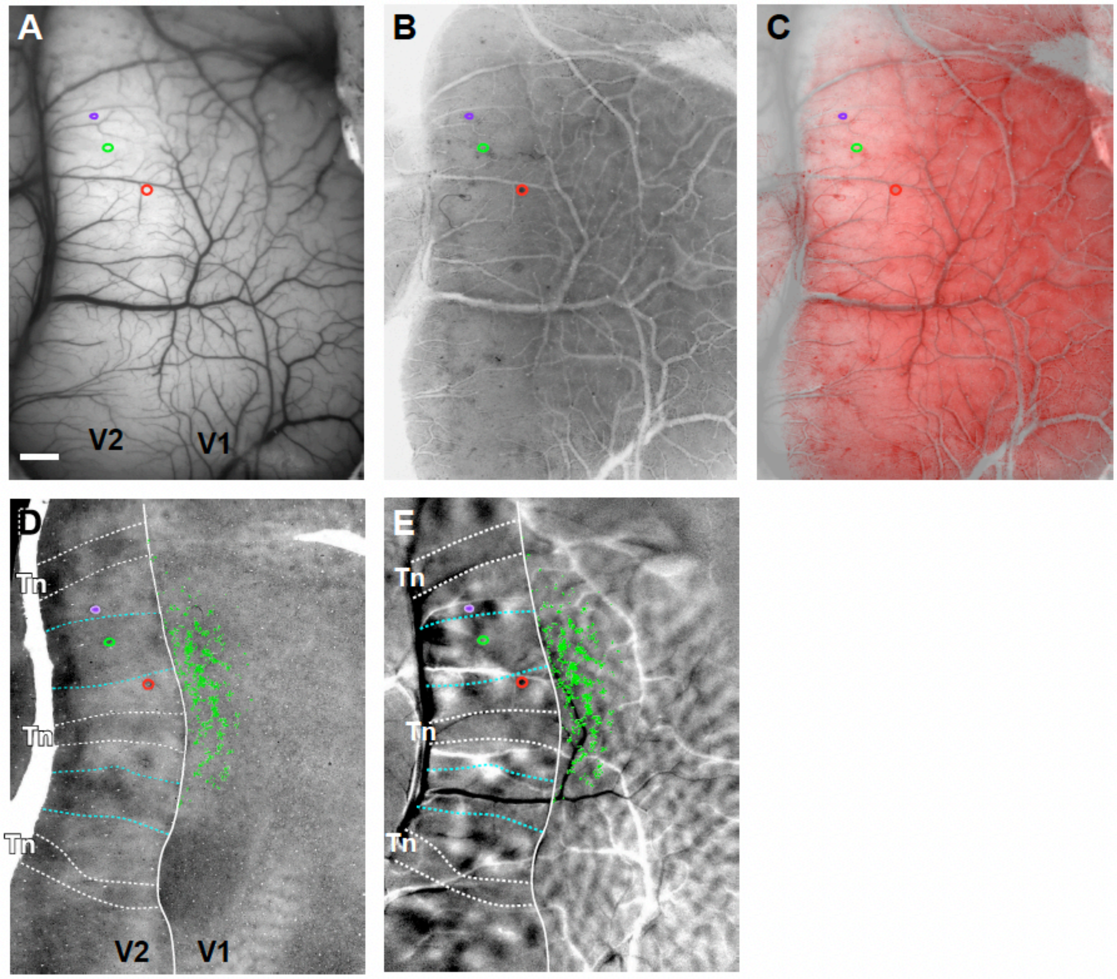


**Extended Data Figure 1. Alignment of histological sections with *in vivo* optical images, and identification of V2 stripe types**

**(A)** *In vivo* image of the cortical surface vasculature in case MK373 (same case as in **Fig. 1**) taken under green light illumination. *Colored* ovals here and in (B-E): outlines of 3 different tracer injection sites. Scale bar : 1mm, valid for all panels. **(B)** The most superficial histological tissue section cut parallel to the imaging plane and stained for CO, showing the surface vasculature running tangentially to the brain surface. **(C)** Overlay of the section in (B) and the optical image in (A) demonstrate excellent alignment of superficial blood vessels. Deeper tissue sections containing cell label or CO stripes are aligned to the superficial section using the radial blood vessels. This approach allows for accurate alignment of labeled cells, injection sites and CO compartments in histological sections to the *in vivo* imaged functional maps. The histological section in (B) was colored red for purpose of illustration. **(D)** A deeper tissue section stained for CO showing the CO stripes in V2. Thin stripes (*Tn*) are outlined in *white*, while thick stripes are outlined in *cyan*. The same stripe outlines are shown superimposed to the orientation maps in panel (E) and in **Fig. 1C-E**. *Green dots:* locations of the CTB488-labeled V1 cells. **(E)** Difference orientation map (same 22.5°-

112.5° map shown in **Fig. 1D**) with superimposed the CO stripe outlines from (D), the 3 tracer injection sites in V2, and the CTB488-labeled V1 cells (*green dots*).

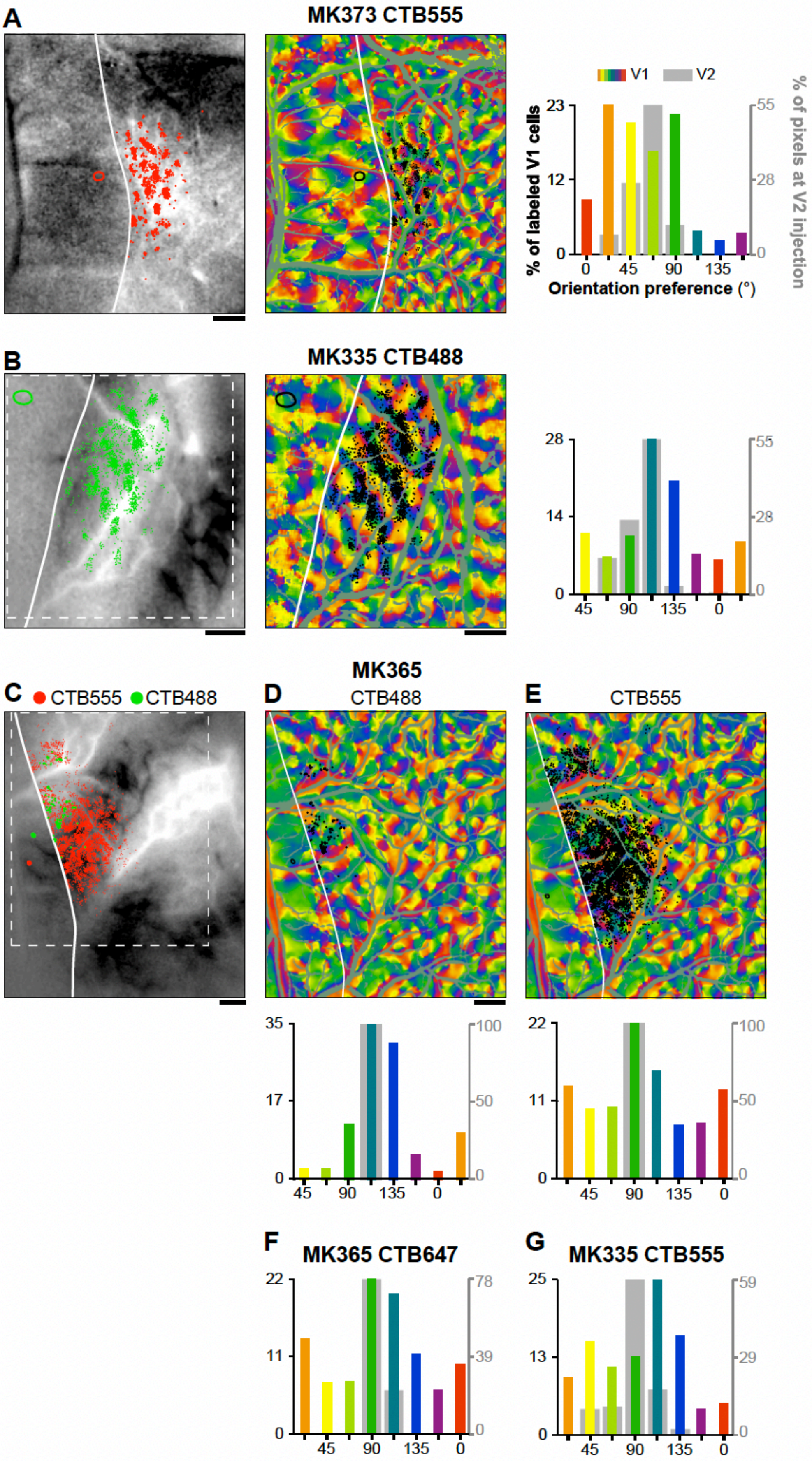

A
MK373 CTB555
V1
V2
% of labeled V1 cells
% of pixels at V2 injection
Orientation preference (°)
B
MK335 CTB488
C
CTB555
CTB488
MK365
D
CTB488
E
CTB555
F
MK365 CTB647
G
MK335 CTB555

**Extended Data Figure 2. Orientation organization of V1 inputs to V2 orientation columns in 6 additional cases**

**(A-E)** Same as panels (A-B) in **Figure 2**, but for 4 different injection cases: MK373-CTB555 (A), MK335-CTB488 (B), MK365-CTB488 (C-D), and MK335-CTB555 (C,E). **(F-G)** For these two additional cases, MK 365-CTB647 (F) and MK335-CTB555 (G) only the orientation histogram of V1 labeled cells are shown.

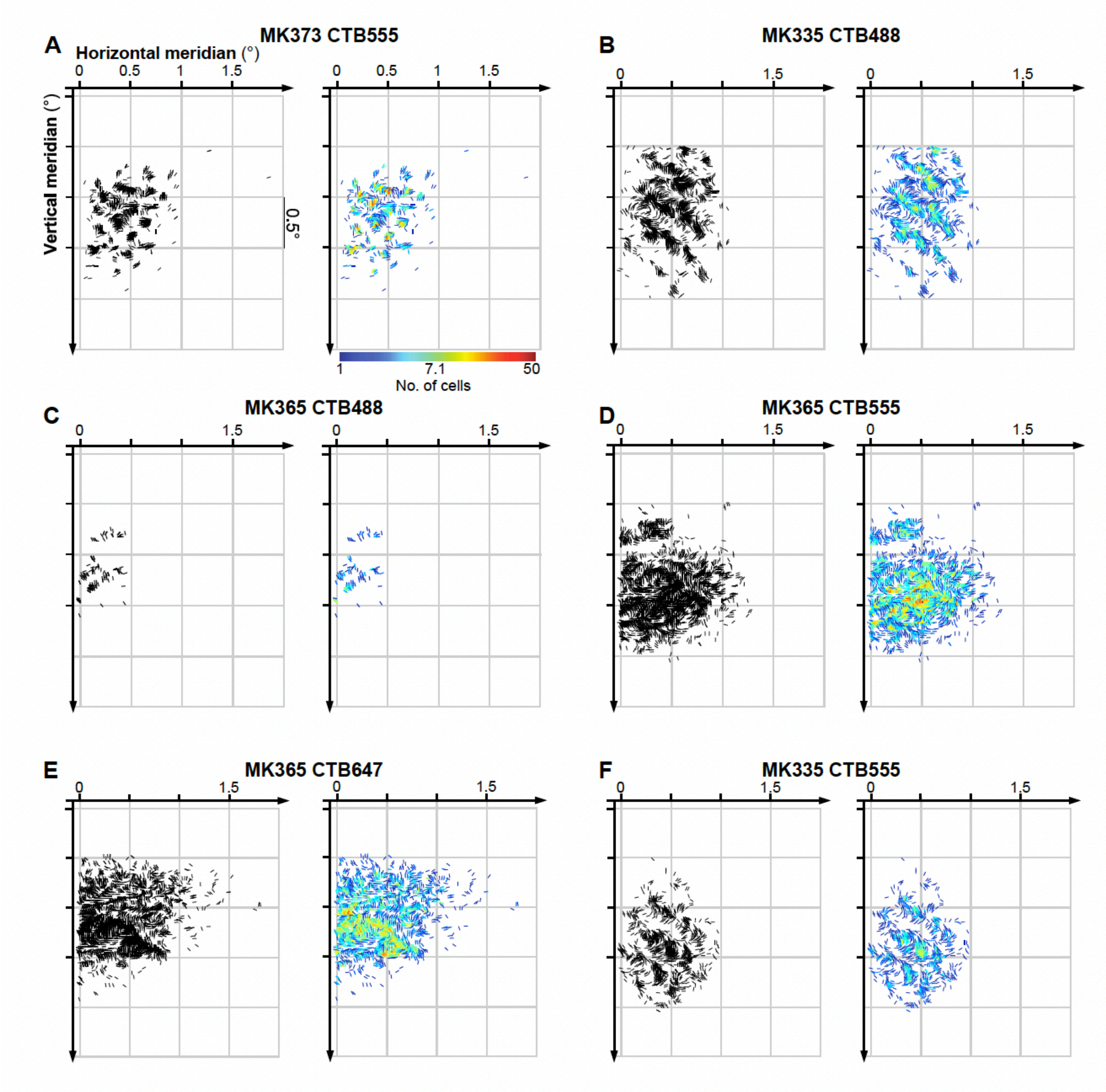


**Extended Data Figure 3. Visuotopic maps of V1 inputs to V2 orientation columns in 6 additional cases**
**(A-F)** Black (LEFT) and color-coded (RIGHT) visual field maps of POs and retinotopic layout of labeled V1 cells for the same 6 injection cases shown in **Extended Data Fig. 2**. Conventions are as in **Fig. 3**.

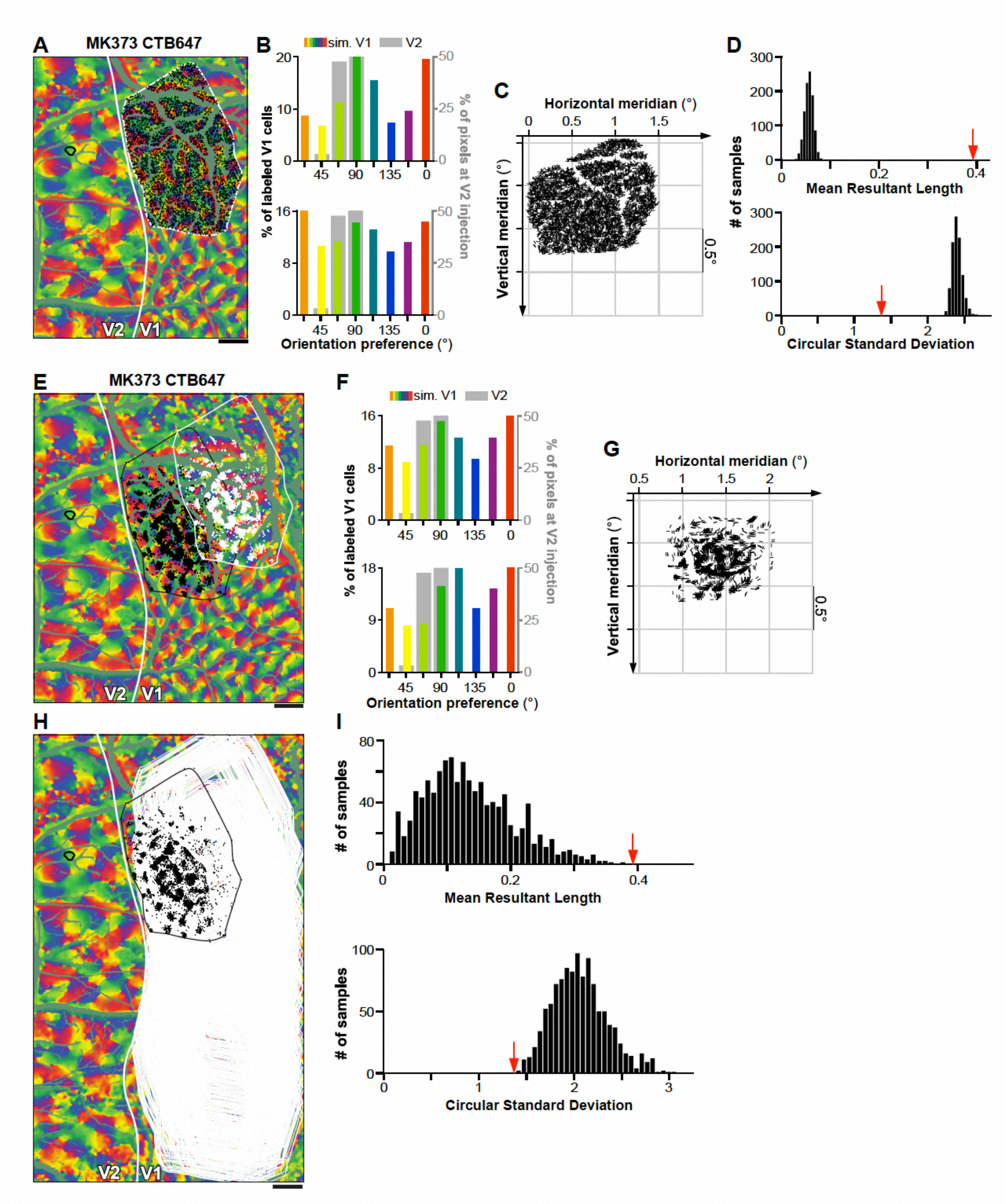


**Extended Data Figure 4. Statistical tests**

**(A)** Control data generated for one example case (MK373-CTB647; same case as in **Fig. 2A-B**). Within the V1 region containing the real labeled cells (*white contour*), we either determined the distribution of POs for all pixels (shown in the top panel of B), or randomly selected as many V1 pixels (*black dots*) as the number of labeled cells in the real data. Other conventions are as in **Fig. 2**. **(B)** TOP: Distribution of POs for all pixels within the white contour in (A). BOTTOM: Distribution of POs for the pixels (*black dots*) selected in (A). *Colored bars:* PO distribution of simulated V1 pixels (*black dots* in A). *Gray bars*: PO distribution at the real V2 injection site. **(C)** The POs of selected pixels for the control V1 data (*black dots* in A) are shown as oriented line segments centered on their corresponding location in visual space. **(D)** Distribution of circular statistics (TOP: mean resultant length; BOTTOM: circular standard deviation) obtained by repeating random pixel sampling from the labeled field 1000 times. The *red arrow* indicates the circular statistics obtained from real data. **(E)** Control data for the same example case generated by shifting the real pattern of cell label from its original location (*black dots*) to a new randomly selected location within the imaged V1 area (*white dots*); in this analysis the relative layout of the real cell label was preserved. **(F)** TOP: Distribution of POs for all pixels within the V1 imaged field of view shown in (E). BOTTOM: Distribution of POs for the control V1 data (*white dots* in E) and for the real V2 data. **(G)** POs and retinotopic mapping for the control V1 data (*white dots* in E and their POs in F bottom). **(H)** The real pattern of cell label (*black dots and contour*) was shifted over the imaged V1 region >1000 times to randomly selected locations (*white contours*) to generate control data. **(I)** Distribution of circular statistics (TOP: mean resultant length; BOTTOM: circular standard deviation) for the control data (*black bars*) obtain by randomly shifting the real pattern of V1 labeled cells. The *red arrow* indicates the circular statistics obtained from real data. Scale bars in (A,E,H): 1 mm.

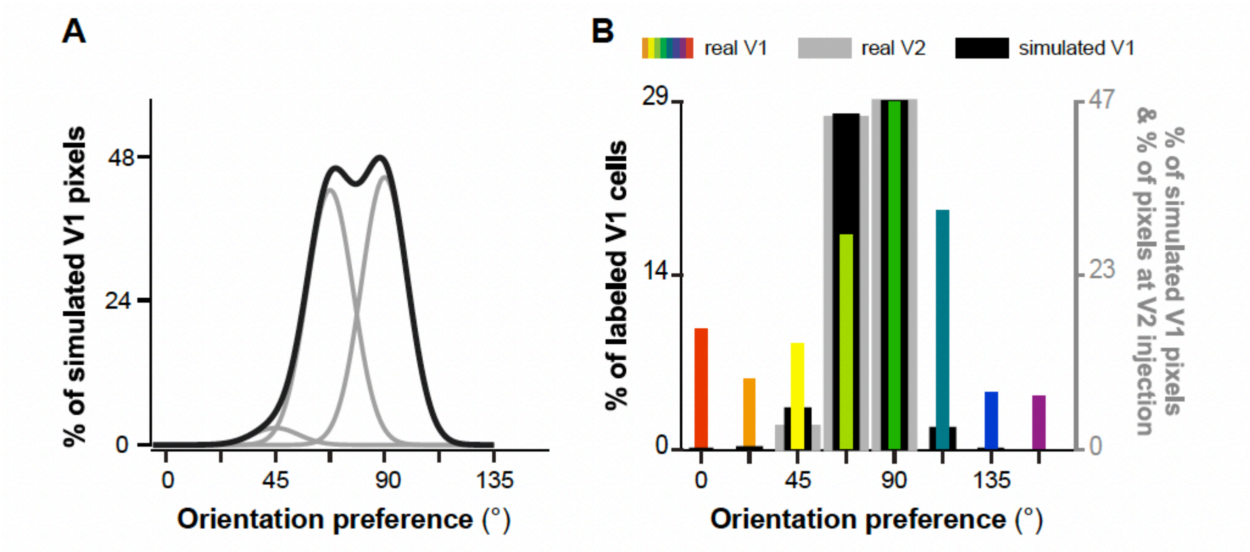


**Extended Data Figure 5. Comparison of PO distribution for real V1-to-V2 connection data with simulated data following a perfect like-to-like connectivity rule.**

**(A)** Distribution of POs of V1 cells predicted by a perfect like-to-like connectivity rule (*black curve*) modeled by weighted summing over three Gaussian functions (*gray curves*) centered at 45°, 67.5°, and 90°, respectively, corresponding to three orientation columns in V2 involved by the real V2 injection site in case MK373-CTB647. **(B)** Simulated PO distribution (after binning) of V1 cells under a perfect like-to-like connectivity rule (*black bars* (see Methods). The simulated distribution differs significantly ($p<0.05$, Chi square comparison) from the distribution of POs obtained from the real V1 data (*colored bars*). *Gray bars*: Distribution of POs at the real V2 injection site.

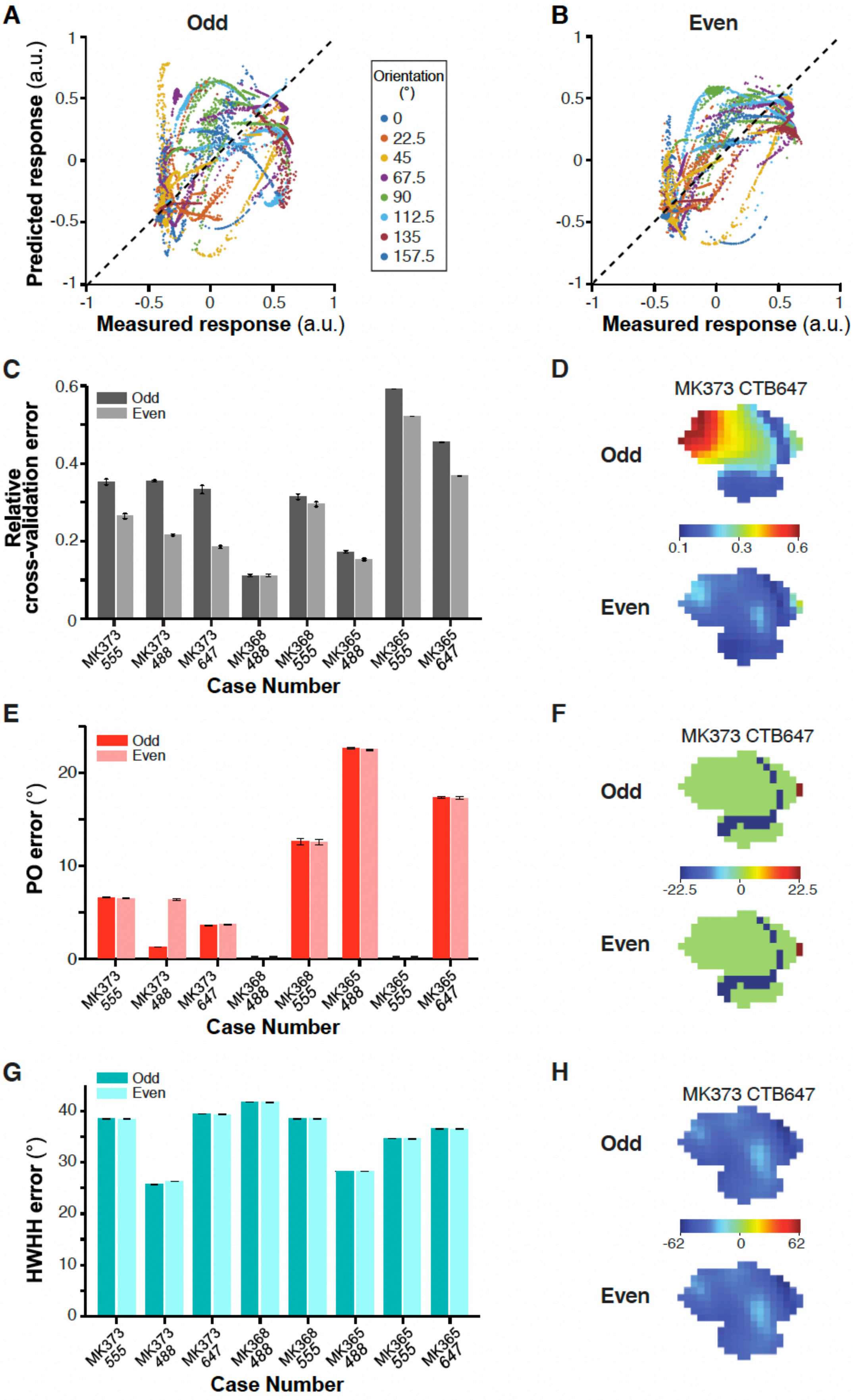

A
Odd
Predicted response (a.u.)
Measured response (a.u.)
Orientation (°)
0
22.5
45
67.5
90
112.5
135
157.5
B
Even
Predicted response (a.u.)
Measured response (a.u.)
C
Relative cross-validation error
Odd
Even
Case Number
MK373 555
MK373 488
MK373 647
MK368 488
MK368 555
MK365 488
MK365 555
MK365 647
D
MK373 CTB647
Odd
Even
0.1
0.3
0.6
E
PO error (°)
Odd
Even
Case Number
F
MK373 CTB647
Odd
Even
-22.5
0
22.5
G
HWHH error (°)
Odd
Even
Case Number
H
MK373 CTB647
Odd
Even
-62
0
62

**Extended Data Figure 6. Performance of the simple-cell feedforward linear model. (A,B)** Responses of V2 cells to excluded grating stimuli of 8 different orientations predicted in the leave-one-out procedure by the two- layer odd and even models, respectively, versus responses measured experimentally. This plot shows data for all the V2 cells/pixels in eight injection cases. The mean correlation coefficients are 0.27+/-0.53 and 0.53±0.44, respectively. **C)** Averaged relative cross-validation error for each injection case , under the odd and even models. (**D**) Color-coded relative cross-validation error maps at the V2 injection site for case MK373-CTB647 under the odd (TOP) and even (BOTTOM) models. **(E)** Averaged absolute error in the model's prediction of the preferred orientation (PO) of V2 cells/pixels under the odd and even models. **(F)** Color-coded maps of signed errors in PO calculated for each pixel at the V2 injection site in case MK373-CTB647 using the odd (TOP) and even (BOTTOM) models. **(G)** Averaged absolute error in the model's prediction of the width of the tuning curves (HWHH) of V2 cells/pixels under the odd and even models. **(H)** Color-coded maps of signed errors in HWHH for V2 pixels in case MK373-CTB647 under the odd (TOP) and even (BOTTOM) models. Error bars: s.e.m.

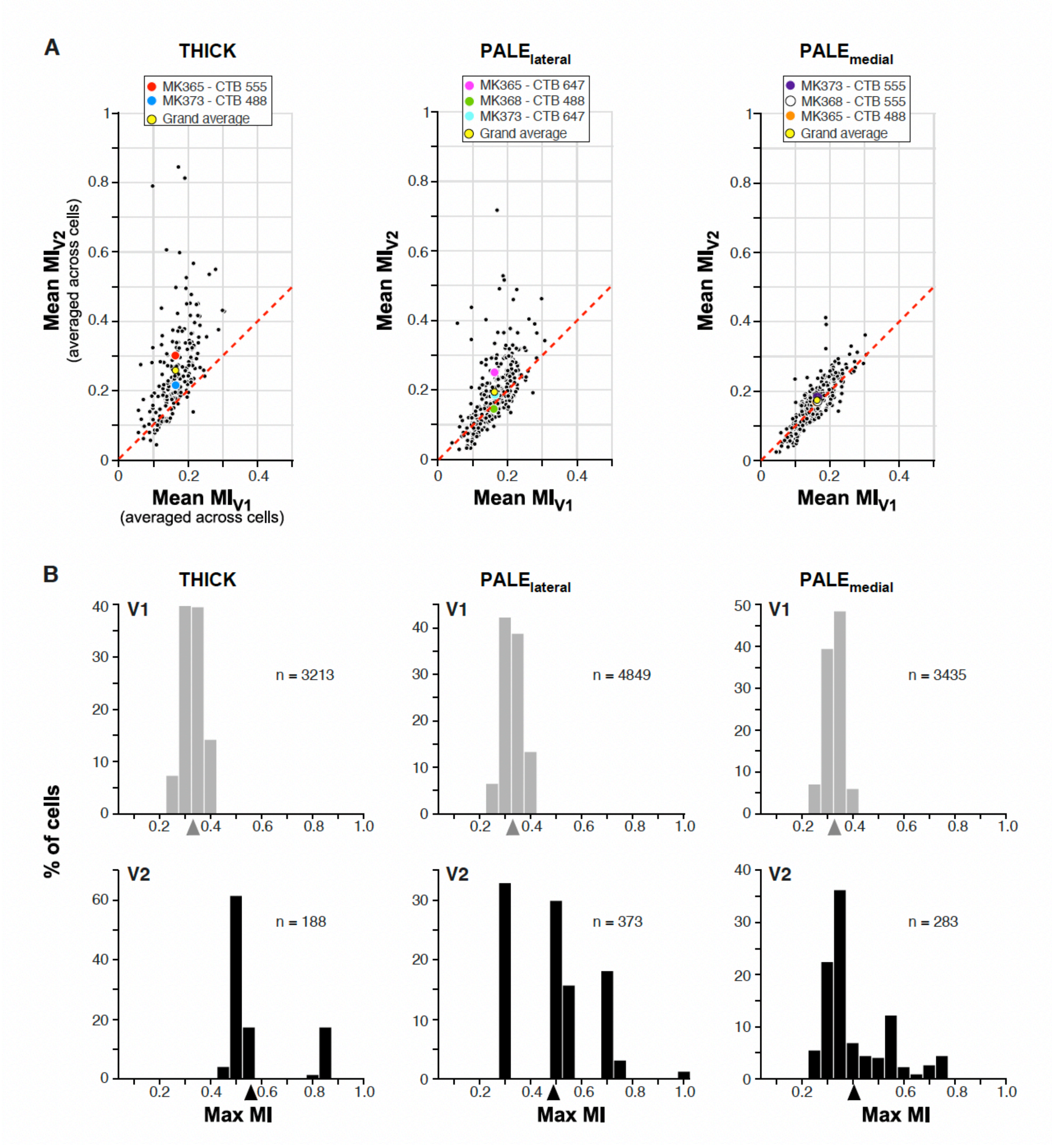


**Extended Data Figure 7. Responses to naturalistic textures of V2 model cells in different V2 CO stripes compared to their V1 input cells.**

**(A)** Each *black dot* in the scatter plots represents the MI for a given texture family averaged across all model V2 cells and their V1 input cells grouped by V2 stripe type. Left: thick stripes; Middle: pale-lateral stripes; Right: pale-medial stripes. *Colored dots* indicate mean MIs across all 97 texture families for each injection case. *Yellow dot* is the grand average across the entire population of cells in that stripe type. **(B)** Distribution of mean MIs for V2 model cells in each stripe type (BOTTOM) and their V1 input cells (TOP). Here, for each V2 and V1 cell we plot the MI with the largest value across all 97 texture families. *Arrowheads*: population mean.

### EXTENDED DATA TABLES

**Extended Data Table 1. Statistical tests for data shown in Extended Data Fig. 4D.** Mean resultant length (MRL) and circular standard deviation (CSD) for real data and control data generated by randomly sampling from the V1 cell labeled field. For each case, we report the range of values for the control data, as well as the fraction of times MRL is larger and CSD smaller than the control data .

| **Case** | **MRL real data** | **Range of MRL for control data** | **CSD real data** | **Range of CSD for control data** |
|---|---|---|---|---|
| **MK373LH - 488** | 0.2<br>F(MRL>0. 2) = 0 | 0.01 – 0.17 | 1.79<br>F(CSD<1.79)= 0 | 1.88 – 3.02 |
| **MK373LH - 555** | 0.56<br>F(MRL>0.56) = 0 | 0.01 – 0.13 | 1.08<br>F(CSD<1.08)= 0 | 2.01 – 2.97 |
| **MK373LH-647A** | 0.32<br>F(MRL>0.32) = 0 | 0.05 – 0.1 | 1.52<br>F(CSD<1.52)= 0 | 2.14 – 2.47 |
| **MK368RH - 488** | 0.6<br>F(MRL>0.6) = 0 | 0.11 – 0.25 | 1.0<br>F(CSD<1.0) =0 | 1.65 – 2.09 |
| **MK365LH - 488** | 0.35<br>F(MRL>0.35) =0.003 | 0.03 – 3.37 | 1.46<br>F (CSD<1.46) = 0 .003 | 1.4 – 2.6 |
| **MK335LH - 488** | 0.27<br>F(MRL>0.27) = 0 | 0.0005 – 0.06 | 1.61<br>FCSD<1.61) = 0 | 2.40 – 3.91 |
| **MK335LH - 555** | 0.22<br>F(MRL>0.22) = 0 | 0.02 – 0. 13 | 1.74<br>F(CSD<1.74) =0 | 2.01 – 2.86 |

**Extended Data Table 2. Statistical tests for data shown in Extended Data Fig. 4I.** Mean resultant length (MRL) and circular standard deviation (CSD) for real data and control data generated by randomly shifting the position of the V1 cell labeled field over the orientation map. For each case, we report the range of values for the control data, as well as the fraction of times MRL is larger and CSD smaller than the control data.

| **Case** | **MRL real data** | **Range of MRL for control data** | **CSD real data** | **Range of CSD for control data** |
|---|---|---|---|---|
| **MK373LH - 488** | 0.2<br>F(MRL>0. 2) = 0.15 | 0.003- 0.38 | 1.79<br>F(CSD<1.79)= 0.15 | 1.38 -3.39 |
| **MK373LH - 555** | 0.56<br>F(MRL>0.56) = 0 | 0.005 – 0.46 | 1.08<br>F(CSD<1.08)= 0 | 1.24 – 3.26 |
| **MK373LH-647A** | 0.32<br>F(MRL>0.32)= 0.018 | 0.003-0.46 | 1.52<br>F(CSD<1.52)=0.018 | 1.24-3.40 |
| **MK368RH - 488** | 0.6<br>F(MRL>0.6)=0 | 0.005 – 0.6 | 1.0<br>F(CSD<1.0) =0 | 1.01 -3.25 |
| **MK365LH - 488** | 0.35<br>F(MRL>0.35) =0.17 | 0.011-0.64 | 1.46<br>F (CSD<1.46) = 0.17 | 0.94- 2.999 |
| **MK335LH - 488** | 0.27<br>F(MRL>0.27) = 0.006 | 0.001 – 0.31 | 1.61<br>FCSD<1.61) = 0.006 | 1.52 – 3.66 |
| **MK335LH - 555** | 0.22<br>F(MRL>0.22) = 0.05 | 0.006 – 0.32 | 1.74<br>F(CSD<1.74) = | 1.51 – 3.83 |
